# Uncertainty-Aware Personalised Gene Expression Prediction Improves Cell-Type-Specific Association Testing

**DOI:** 10.64898/2026.11.06.764604

**Authors:** Patrick Styll, Yazheng Di, Na Cai

**Affiliations:** Department of Biosystems Science and Engineering, ETH Zurich, Basel, Switzerland; Department of Computer Science, ETH Zurich, Zurich, Switzerland

## Abstract

Linking genetic variation to disease requires accurate prediction of its effects on gene expression in relevant cell types. Sequence-to-function models offer a way to predict these effects and, with training on personal genomes, have begun to capture inter-individual expression differences. However, effect sign accuracy remains limited, and predictive uncertainty is unquantified. Here we introduce ATTICUS, which combines personalised sequence representations with uncertainty-aware expression prediction. ATTICUS transfers regulatory information learned in bulk expression pretraining to cell-type-specific expression prediction without requiring genotypes paired with single-cell expression data. Across blood and brain cell types in OneK1K and ROSMAP, ATTICUS approximately doubles expression prediction correlations relative to scPrediXcan and substantially improves sign accuracy. Moreover, ATTICUS enables uncertainty-aware transcriptome-wide association studies (TWAS). Applied to systemic lupus erythematosus and Alzheimer’s disease, ATTICUS identifies more associations, with greater cell-type specificity and stronger directional concordance, and reduced statistical inflation. Together, our results demonstrate that personalised training makes predictions more directionally consistent, and uncertainty propagation improves the calibration of TWAS and the prioritisation of disease genes. ATTICUS provides a framework for both, enabling the use of single-cell expression datasets without matched genotypes to inform uncertainty-aware TWAS, expanding their use in prioritising disease-associated genes and cell types.

## 1 Introduction

Genetic variation shapes gene expression differently across cell-types, and recent studies have converged on the revelation that cell-type-specific gene regulatory effects are most likely central to understanding complex traits and diseases [1, 2, 3, 4]. Single-cell transcriptomic atlases now span millions of cells and hundreds of cell-types [5, 6, 7], many of which profiled in tens to hundreds of people. Genotyped cohorts large enough to map genetic variant effects on these transcriptomic profiles however exist only for relatively few tissues, chiefly peripheral blood [8] and the cerebral cortex [9, 10, 11, 12, 13]. Genetic effects on expression therefore remain uncharacterised in most catalogued cell-types, and cannot be combined with genome-wide association study (GWAS) findings in transcriptome wide association studies (TWAS) [14, 15] to determine the genes and cell-types relevant to complex traits and diseases.

Sequence-to-function (S2F) models such as Enformer [16], Borzoi [17], and AlphaGenome [18] learn the regulatory relationship between genetic sequence and gene expression levels in bulk-tissues, and offer the opportunity to transfer them to predictions of cell-type level gene expression through further training. Several methods have now attempted transferring the S2F learnt regulatory effects to predicting single-cell or cell-type resolved gene expression (e.g., UNICORN [19], seq2cells [20], scPrediXcan [21], Decima [22], scooby [23]). However, being trained on the human reference genome, these S2Fs seem unable to capture how variations across individuals affect gene expression, often yielding weak or directionally inconsistent predictions across individuals [24, 25, 26, 27, 28, 29]. Personalised S2F models, trained on paired genome–transcriptome data across individuals [30, 31, 32, 33], have recently been developed to address this limitation [34, 35, 26, 36, 37]. Among them, VariantFormer [36] was trained on 21,004 samples from 2,330 donors across multiple tissues, and predicts expression differences across donors comparably to genotype-based models and better than reference-trained S2F models. Whether personalised pretraining transfers from bulk tissues to cell-types, and whether it yields directionally correct predictions across individuals at the cell-type level, remains underexplored [38].

When it comes to applying learnt regulatory effects and their predicted cell-type level gene expressions for downstream applications like TWAS, the ideal model should estimate, propagate and utilise meaningful uncertainty measures to produce robust findings [39]. Few S2F models come with uncertainty estimates [40, 19], and most have been shown to give predictions with varying reliability [24, 25, 41, 21, 29]. Likewise, cell-type resolved TWAS applications of S2F models like ctPred in scPrediXcan [21] return point estimates of cell-type level gene expression and their associations with phenotypes. This reveals a gap: treating S2F predicted expression as error-free understates the standard errors of TWAS statistics and incurs false-positive findings [42, 43]; existing methods that propagate uncertainty account only for estimation uncertainty in SNP weights derived from linear models and were developed only for TWAS in bulk tissues [44, 43, 45, 46, 47]. To our knowledge, no method propagates uncertainty in a nonlinear S2F predictor or has been applied to single-cell TWAS.

We therefore developed ATTICUS (**a**llele-aware **t**ranscript **t**rait **i**nference in single **c**ells with **u**ncertainty-aware **s**tatistics), a framework for personalised cell-type specific gene expression prediction from S2F models that enables uncertainty-aware TWAS. ATTICUS enables the transfer of regulatory information from variant-aware S2F pretrained on bulk genome–transcriptome cohorts to cell-type specific expression prediction, through training on S2F embeddings with single-cell gene expression data without requiring matched genotypes. We demonstrate that ATTICUS achieves higher accuracy in cell-type level gene expression prediction across genes and across cells in both blood cell-types in OneK1K [8] and brain cell-types in ROSMAP [9, 10, 11] than current state-of-the-art models. Most importantly, when paired with a variant-aware S2F like VariantFormer [36], ATTICUS achieves high sign accuracy in personalised cell-type level gene expression predictions, a substantial improvement from near-random sign accuracies shown in previous benchmarks [24, 25]. ATTICUS is also designed so that it can account for uncertainty from expression prediction through association testing: it models variance in single-cell gene expression inputs, estimates uncertainty in the fitted cell-type specific predictors using a deep ensemble [48] training architecture, and preserves uncertainty estimates in the SNP weights used for downstream TWAS analyses through a probabilistic distillation process. We demonstrate that uncertainty-aware TWAS using ATTICUS’ probabilistic distillation on heterozygote-aware VariantFormer predictions gives higher numbers of better-calibrated and disease-specific TWAS hits when applied to systemic lupus erythematosus (SLE) and Alzheimer’s disease (AD) than existing approaches.

## 2 Methods

### ATTICUS Overview

We introduce ATTICUS, a framework that enables prediction of personalised cell-type specific gene expression. ATTICUS transfers regulatory information learnt from bulk genome-transcriptome cohorts in S2F models to single-cell RNA-seq data without linked genotype data. It takes as input cis-regulatory context embeddings from the variant-aware pretrained S2F model Variant-Former [36], as well as cell-type-specific pseudobulk gene expression data from scRNA-seq cohorts like OneK1K [8] and ROSMAP [9, 10, 12]. This framework consists of three stages: a) uncertainty-aware personalised gene expression prediction, b) probabilistic distillation into genotype-based linear predictors, and c) propagation of the resulting distribution over expression models into TWAS association statistics (**Figure 1**).

**Figure 1:**
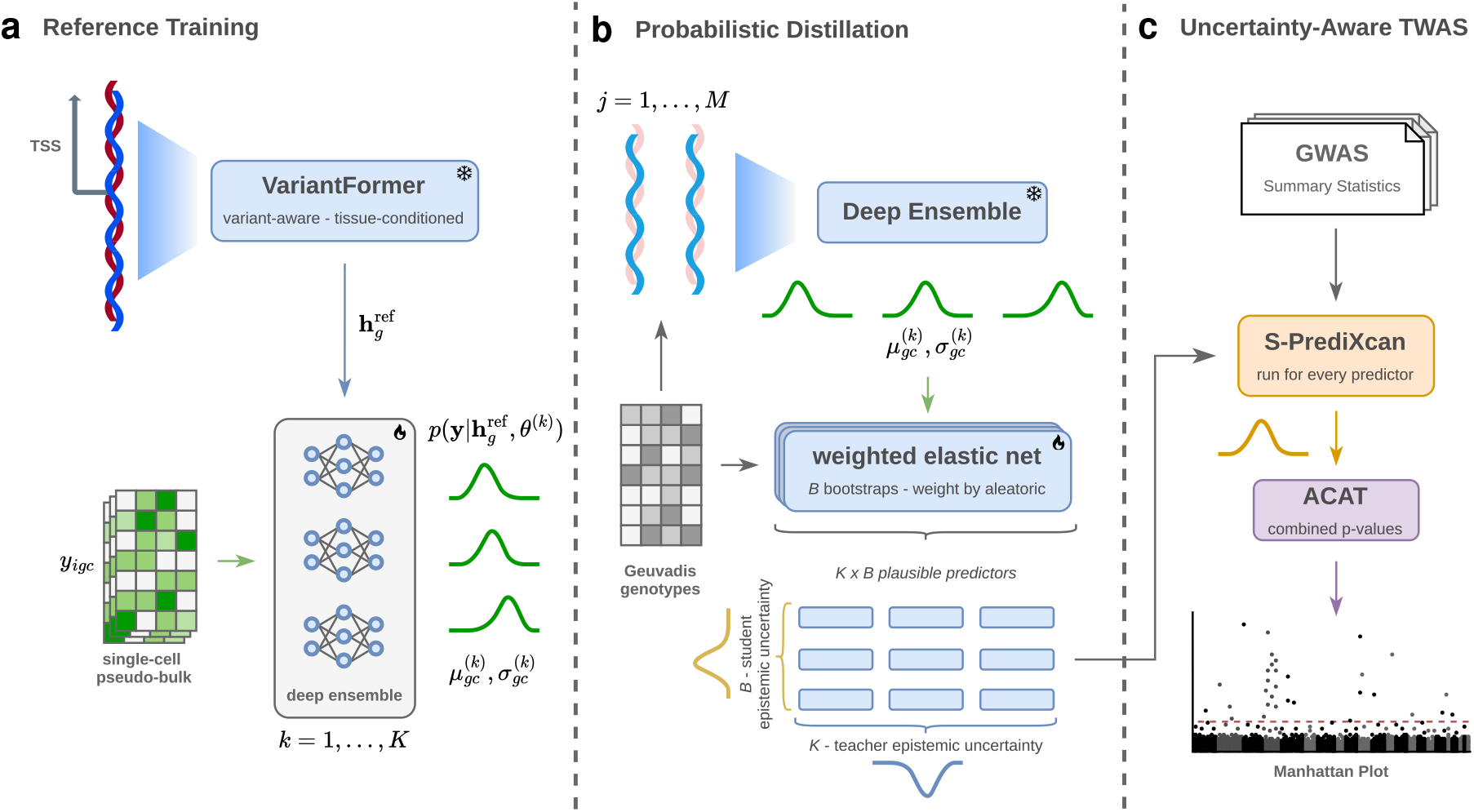
Overview of the ATTICUS framework. a) ATTICUS is a deep ensemble based on an S2F backbone, trained to predict cell-type specific pseudobulk expression, which is then b) distilled into genotype-based linear predictors and c) used for uncertainty aware TWAS.

### 2.1 Uncertainty-aware personalised gene expression prediction

#### Pairing variant-aware, bulk-tissue S2F representations with pseudobulk gene expression

For each individual *i* = 1, … , *N* , gene *g* = 1, … , *G*, and cell-type *c* = 1, … , *C* from scRNA-seq cohorts we use in our model training in OneK1K [8] and ROSMAP [9, 10, 12], let *y*_*igc*_ ∈ ℝ denote the observed pseudobulk gene expression. Notably, we model pseudobulk gene expression levels as quantitative expression, rather than as a relative rank across genes per cell-type as performed in scPrediXcan [21], to preserve the interpretability of our predictions as expression magnitudes. We split individuals in these cohorts into an 80% training-individuals set and a 20% test-individuals set. For genes in the trainingindividuals, we split them by chromosomes into training-genes, validation-genes, and test-genes sets in the same way as previously done in other methods for ease of benchmarking (**Appendix Note B.2**).

For each training-gene *g* in each training-individual we obtain a fixed S2F embedding 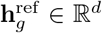 on the 1 MB GRCh38 human reference sequence cis-regulatory context surrounding its transcription start site (TSS). For the S2F backbone, we use VariantFormer as a tissue-conditioned backbone for representations of personalised genetic effects. This embedding is shared across all training-individuals used for *teacher training* (**Appendix Note B.2**): the model observes the same sequence representation 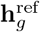 paired with individual and cell-type-specific expression measurements 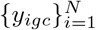. Pairing a common reference-sequence S2F representation with pseudobulk gene expression from multiple training-individuals enables the teacher to recover both the population-average and the individual variability in gene expression levels per cell-type (**Appendix Note C**).

#### Probabilistic training

To capture uncertainty in the fitted predictor, we represent the teacher as a deep ensemble of *K* independently initialised probabilistic neural networks, where we denote the parameters of ensemble member *k* = 1, … , *K* as 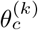. For each cell-type *c*, we learn a probabilistic predictor 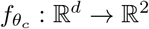 , with

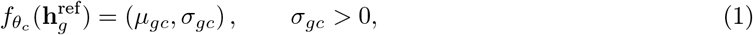

where *µ*_*gc*_ and *σ*_*gc*_ denote the conditional mean and standard deviation of expression for gene *g* in cell-type *c* (the latter is referred to as *aleatoric* uncertainty). The ensemble predictive mean is obtained by averaging the predicted means across members. Total *aleatoric uncertainty* is given by the average variance predicted by the individual ensemble members, which we find to be representative of population-wide expression variance (**Appendix Note C**); *epistemic uncertainty* is given by the disagreement between their predicted means (**Appendix Figure 1**).

Each ensemble member is trained using a learnt weighted combination of Pearson correlation, cosine similarity, Poisson negative log-likelihood (PNLL) and Gaussian negative log-likelihood (GNLL) losses (**Appendix Note B**). The GNLL term explicitly controls the predicted variance, which enables the model to learn aleatoric uncertainty alongside expression predictions. We select hyperparameters based on the predictions in the validation-genes across training-individuals.

#### Teacher evaluation

We first assessed teacher prediction performance across test-genes per cell-type, and cell-types per test-gene across all training-individuals, through a) fixing a cell-type and computing the Pearson correlation across test-genes and b) fixing a test-gene and computing the Pearson correlation across cell-types. Correlation of predicted and observed gene expression levels of a single gene across cell-types is a harder task given that such variation is lower than variation across genes, and measures how well the model can predict cell-type-specific gene expression. We then evaluated personalised predictions across test-individuals across all genes (including training-, validation- and test-genes) and cell-types, restricting evaluations to cell-types represented across all test-donors (**Appendix Table 1**).

### 2.2 Probabilistic distillation

To obtain genotype-based predictor weights suitable for TWAS, we distill the nonlinear teacher ensemble into linear (student) models. Given genotype data of *M* individuals (not necessarily used in teacher training), we construct the personalised sequence of each individual *j* = 1, … , *M* around gene *g* and obtain the corresponding S2F embedding 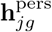. Each teacher ensemble member provides personalised pseudo-expression predictions, yielding a panel of paired genotype–pseudo-expression data:

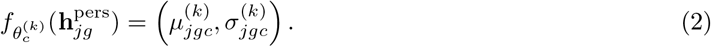

For each gene *g*, let **x**_*jg*_ denote the genotype dosages of the cis-variants used by the student model for individual *j*. For each cell-type *c* and teacher ensemble member *k*, we fit a separate elastic-net predictor

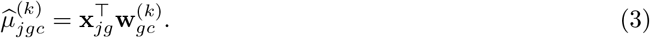

to approximate the corresponding personalised teacher predictions 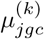.

Not all teacher predictions are equally informative for TWAS. As such, we down-weigh observations with higher *aleatoric uncertainty*, while up-weighting those with lower uncertainty, as previously demonstrated in scTWAS [49]. The student weights are estimated by minimizing the uncertainty-weighted elastic-net objective

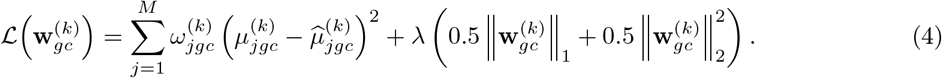

where 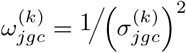.

We then quantify *student uncertainty* using bootstrap resampling of the *M* individuals used for distillation, as previously demonstrated in BAY-TS [44]. For each teacher ensemble member *k* and bootstrap replicate *b* = 1, … , *B*, we independently refit the uncertainty-weighted elastic net, yielding 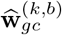 ; disagreement in the fitted SNP weights across bootstrap replicates *b* of the same teacher member *k* captures *student uncertainty*. Together, the *K* × *B* fitted plausible predictors define an empirical distribution over genotype-based expression models, which we propagate into downstream TWAS.

Unlike Enformer, which is used by scPrediXcan and processes a single DNA sequence, VariantFormer explicitly represents diploid genotypes [36], allowing heterozygote-specific prediction effects that a dosage-only additive student cannot fully capture. We therefore jointly fitted additive SNP-dosage and heterozygosity coefficients using the uncertainty-weighted elastic-net objective (**Eq. 4**). To remain compatible with S-PrediXcan and additive GWAS summary statistics [50], we subsequently project these coefficients into effective additive weights [51] (**Appendix Note B.3**).

### 2.3 Propagating uncertainty into S-TWAS

Conventional summary-statistic TWAS (S-TWAS) uses a single set of weights on a single set of predictors. In contrast, our probabilistic distillation yields an empirical distribution of *K* × *B* plausible genotype-based predictors and their weights, reflecting both teacher and student uncertainty. We ran S-PrediXcan separately for each of the *K* × *B* fitted student models; for each gene–cell-type pair, this produces *K* × *B* TWAS *Z*-statistics, effect estimates, and *p*-values. These values form empirical distributions that describe how the association results vary across plausible expression models under both teacher and student uncertainty. We then combined the non-independent model-specific *p*-values using the aggregated Cauchy association test (ACAT) [52, 53].

## 3 Results

### 3.1 Quantitative expression targets and tissue-conditioned S2F representations improve reference prediction

We first evaluate how well ATTICUS predicts gene expression across all test-genes and cell-types in held-out genes in the 80% of training-individuals in OneK1K and ROSMAP (**Figure 2a, Appendix Note B.2**), using correlations between predicted and observed values, and compare its performance to ctPred implemented in scPrediXcan. We expect achieving a high across cell-types prediction performance is the harder task given that variation of the expression of a single gene across cell-types is lower than that across genes, and measures how well the model can predict cell-type-specific gene expression. The latter task is closer to what S2Fs are trained to do - learn transferrable regulatory grammar across genes within a bulk tissue - and the performance measures how this transfers to cell-type-specific gene expression.

**Figure 2:**
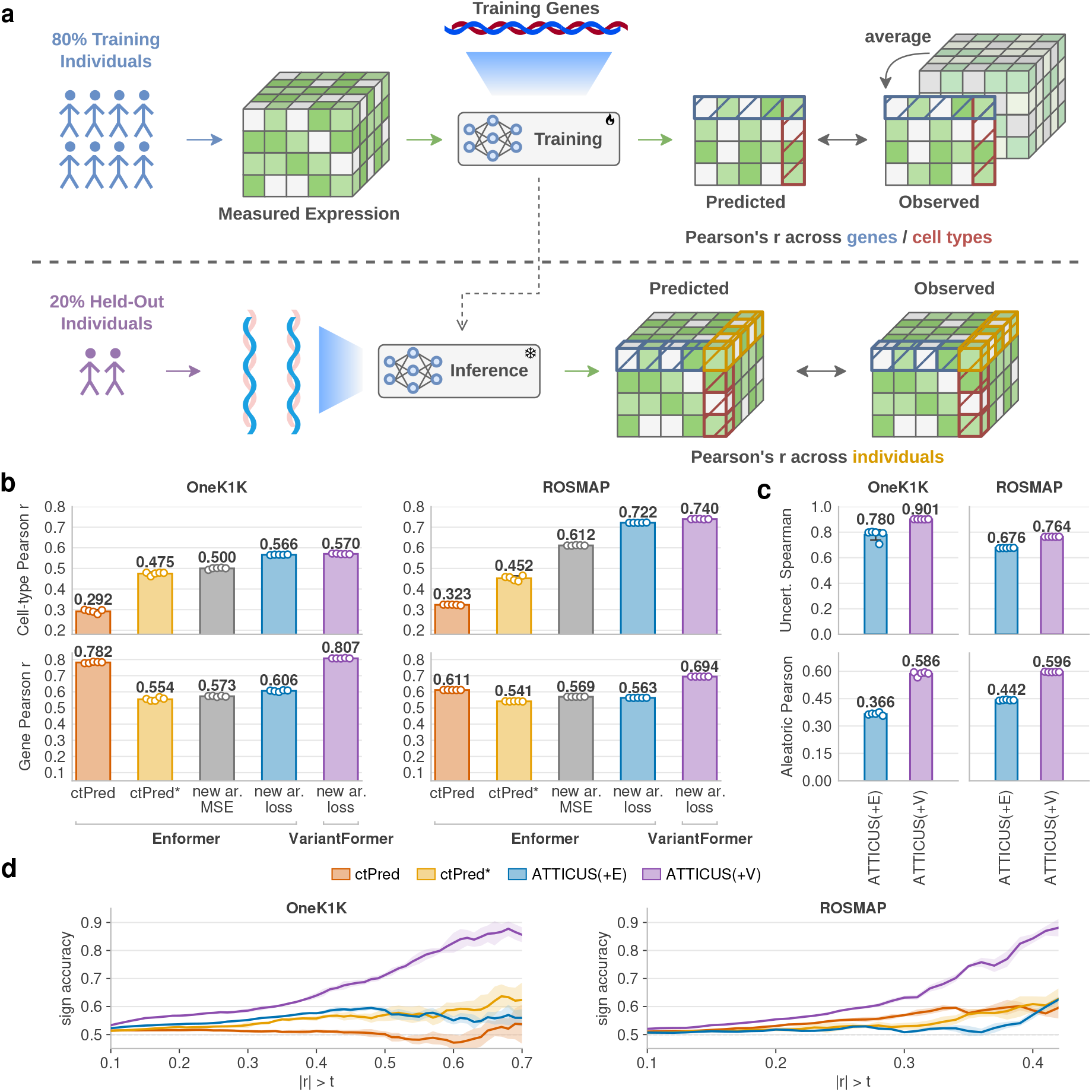
Benchmarking predictive performance. a) Overview of evaluation in reference- and personalized prediction. b) Reference-genome prediction on held-out chromosomes. c) Correlation of epistemic uncertainty with prediction error (Spearman) and aleatoric uncertainty with population-wide expression variance (Pearson). d) Personalized prediction in held-out individuals. The x-axis shows thresholds t on the absolute correlation between predicted and observed expression across individuals. Beware that estimates at high thresholds are based on progressively fewer gene–cell-type pairs.

Our full ATTICUS implementation with variant-aware, tissue-conditioned VariantFormer S2F representations yields the best predictions on both levels, achieving across-gene correlations of 0.807 and 0.694 for OneK1K and ROSMAP respectively, and across-cell-type correlations of 0.57 and 0.74 for the two cohorts (**Figure 2b**). In comparison, ctPred implemented exactly as in scPrediXcan, using the Enformer S2F backbone, gives similar across-gene correlations of 0.782 and 0.611 for OneK1K and ROSMAP, respectively, but substantially lower across-cell-type correlations of 0.292 and 0.323. In other words, our full ATTICUS implementation performs substantially better than SOTA on both tasks. We therefore set out to understand how much each component of our model contributes to this improved performance.

First and most notably, ctPred ranks log-transformed pseudo-bulk expression levels of genes per cell-type and then converted into percentiles ranging from 0 to 1. It then predicts gene-expression percentiles, rather than the log-transformed pseudo-bulk. This is a different prediction target than ATTICUS, and likely explains why ctPred does well at across-gene predictions within a cell-type, but poorly in across-cell-type predictions. We therefore train a second version of ctPred, denoted ctPred^*^, directly on log-transformed pseudo-bulk expression like we do in ATTICUS. This model achieves higher across-cell-type correlations of 0.475 and 0.452, alongside lower across-gene correlations of 0.554 and 0.541, respectively. The former confirms our hypothesis, and the latter demonstrates that the task ATTICUS focuses on, predicting expression on its quantitative scale, is the more difficult one that requires changes to both the neural network architecture and loss function (**Appendix Note B.2**).

We therefore ask whether the deep ensemble architecture and composite loss functions in ATTICUS contribute to improvement in prediction performance, especially in terms of learning cell-type specific expression regulation. To do this, we first replace the ctPred* architecture with our deep ensemble architecture, and found that it increases across cell-type correlations from 0.475 to 0.500 in OneK1K and from 0.452 to 0.612 in ROSMAP. Across-gene correlations also improve slightly. We think this is because the uncertainty estimates we derive from the deep ensemble capture information on prediction reliability and inter-individual expression variability (**Figure 2c**) and generalise to unseen genes, in a way that improves with S2F model performance. Second, we replace the MSE loss implemented in ctPred* with our composite loss (**Appendix Note B.2.4**), which further improves across-cell-type correlations to 0.566 and 0.722, while across-gene correlations reach 0.606 and 0.563 (**Figure 2b**). Both architectural modifications and composite loss in ATTICUS result in substantial improvements in recovering cell-type-specific expression patterns, especially in ROSMAP. Overall, we find improvements in cell-type specificity of predicted gene expression levels are well-accounted for by the deep ensemble architecture and composite loss functions in ATTICUS. We do not attribute this gain to the S2F backbone choice of VariantFormer over Enformer.

We attribute the remaining gap in performance of ctPred* trained with our full architecture and composite loss, relative to our full ATTICUS model, to the difference in S2F representations in VariantFormer (in ATTICUS) and Enformer (in scPrediXcan). Apart from having been trained in a variant-aware manner with thousands of diploid genomes rather than just the human reference genome, VariantFormer uses a different sequence context than Enformer (**Appendix Note B.2.1**). Detailed ablation studies on these models are needed to assess how each aspect of sequence context and variant-aware training contribute to their improved regulatory context representation - we think of this as out of scope for this paper. For the rest of this paper, we refer to our full ATTICUS implementation with VariantFormer as ATTICUS(+V), and the benchmark it against the ATTICUS implementation with Enformer, ATTICUS(+E), as well as ctPred in scPrediXcan.

### 3.2 Personalized bulk pretraining improves the directionality of expression prediction across donors

We evaluate personalised prediction on all genes in the 20% held-out test-individuals in OneK1K and ROSMAP (200 and 81 individuals, respectively, **Figure 2a**), restricting comparisons to cell-types represented across all test donors (**Appendix Table 1**). When comparing the performance of our full model ATTICUS(+V) with ATTICUS(+E) and ctPred in scPrediXcan, we note three important considerations: first, ctPred predicts percentiles of gene expression within cell-types, while our ATTICUS models predict log-transformed pseudo-bulk expression; second, we evaluate prediction accuracy using the mean absolute across-individual Pearson correlation between predictions and observed values, rather than the *m*_1_ statistic [54] used by scPrediXcan (**Appendix Note D, Appendix Figure 2**); third, in addition to evaluating mean absolute across-individual Pearson correlation, we further evaluate the sign accuracy of our predictions, defined as the fraction of positive correlations among genes satisfying | *r*| *> t*, where *t* is a threshold on absolute across-individual Pearson correlation.

We find that all models perform well across genes and cell-types, but generally badly across test-individuals (**Appendix Figure 3a**,**b**). Using VariantFormer rather than Enformer as the S2F back-bone in ATTICUS does not improve mean absolute across-individual Pearson correlation, despite its stronger reference-prediction performance. This can be partially accounted for by aleatoric and epistemic uncertainty estimates in ATTICUS (**Methods, Appendix Figure 4**). Nevertheless, strikingly, ATTICUS(+V) substantially improved across-individual prediction sign accuracy in both OneK1K and ROSMAP (**Figure 2d**): at | *r*| *> t* = 0.4, ATTICUS(+V) sign accuracy reaches approximately 65% in OneK1K and 80% in ROSMAP, whereas the Enformer-based approaches including ATTICUS(+E) remained around guessing levels. Most importantly, sign accuracy of ATTICUS(+V) increases with mean absolute across-individual Pearson correlation (**Figure 2d**). Restricting our evaluation to genes with high cis-predictability improved both absolute correlations and sign accuracy further (**Appendix Figure 5**). Overall, we demonstrate, for the first time, improved sign accuracy in personalized gene expression prediction beyond guessing levels [24, 25].

**Figure 3:**
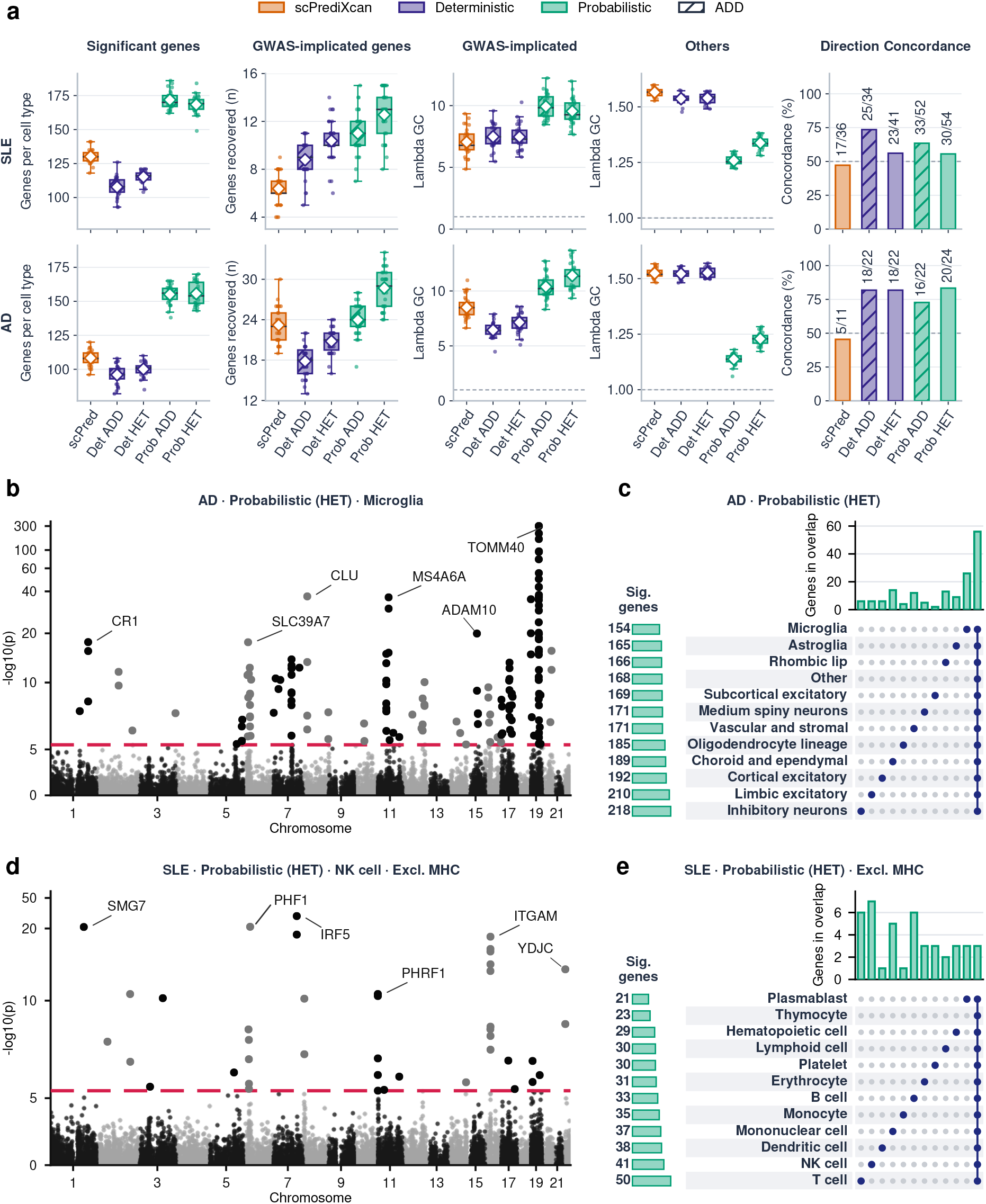
Uncertainty-aware TWAS improves recovery of disease-associated genes. a) Comparison of scPrediXcan with deterministic and probabilistic ATTICUS(+V) using additive (ADD) and heterozygote-aware (HET) models. b,d) Manhattan plots of probabilistic ATTICUS(+V) HET S-TWAS associations in AD microglia and SLE NK cells, respectively. c,e) UpSet plots showing significant genes across 12 major cell-type groups in AD and SLE, respectively, using probabilistic ATTICUS(+V) HET S-TWAS associated hits.

We then seek to investigate if this improvement in sign accuracy can be attributed entirely to variant-aware pretraining. To do this, we assess ATTICUS predictions across individuals using another S2F model, Variformer [34], which fine-tunes Enformer on paired personal genomes and bulk expression while retaining its architecture. We refer to this as ATTICUS(+VE). On the approximately 300 genes used for its Whole Blood fine-tuning, both ATTICUS(+VE) and ATTICUS(+V) substantially improve sign accuracy relative to ATTICUS(+E), with comparatively modest changes in absolute correlations (**Appendix Note E**; **Appendix Figure 6b**,**c**). These results suggest that improved sign accuracy mainly arises from variant-aware pretraining, while gap between ATTICUS(+VE) and ATTICUS(+V) may reflect better representations in VariantFormer learnt through both variant-aware and tissue-conditioned training at scale and across all genes.

### 3.3 S2F models pretrained on bulk tissue do not capture cell-type specific sequence regulatory effects

An important limitation previously not explicitly demonstrated is that all S2F models trained on bulk tissue gene expression may fail to capture cell-type specific sequence regulatory effects, particularly those that are inconsistent with bulk tissue. To demonstrate this, we stratify gene–cell-type pairs according to whether observed inter-individual expression differences are consistent or inconsistent with directions predicted by zero-shot VariantFormer in bulk tissue. Overall, sign accuracy in ATTICUS(+V) was substantially higher when the observed directions are consistent with the bulk prediction, whereas effects inconsistent with it remained difficult to recover (**Appendix Figure 7**).

We then asked if sign accuracy in ATTICUS(+V) is only improved in genes with high cell-type shared cis-heritability, estimated by CIGMA [4], and not in genes with high cell-type specific cis-heritability. Our findings are exactly consistent with our hypothesis. While ATTICUS(+V) performs better in terms of prediction and sign accuracy than other models across a range of cell-type shared and cell-type specific cis-heritability, this gain is much more prominent among genes with high cell-type shared heritability (**Appendix Figure 8**). We believe this is because VariantFormer, the S2F ATTICUS(+V) uses, is only trained on bulk tissues. Future models trained directly on single-cell gene expression should resolve cell-specific effect directions.

### 3.4 Probabilistic and heterozygosity-aware distillation preserves uncertainty for down-stream analyses

To enable summary-statistic TWAS, we distill the nonlinear predictors into genotype-based elastic-nets via deterministic and probabilistic versions of ATTICUS(+V) using genotypes of 503 individuals in the 1000G Phase 3 European dataset [55]. We train the distillation model in 80% of these individuals, and evaluate distillation fidelity using gene-wise teacher-student Spearman correlations across the 20% held-out individuals (**Appendix Note B.3**).

Notably, ATTICUS(+V) enables heterozygote-ware (HET) distillation (**Methods**, **Appendix Note B.3**): both deterministic and probabilistic HET distillation in ATTICUS(+V) achieve high distillation fidelity (mean correlations of 0.77 for both cell types in OneK1K and ROSMAP) similar to Enformer-based l-ctPred and l-ctPred^*^ models (**Appendix Figure 9a, Appendix Table 2**). Probabilistic distillation, in particular, enables propagating teacher and student uncertainty into downstream association testing. We also find that while projecting the heterozygosity-aware students (HET pure) onto additive predictors (HET) reduces their fidelity to the teacher (**Appendix Figure 9b**), they remain higher in distillation fidelity than heterozygosity-nonaware students trained directly using SNP dosages alone (ADD). It is open to investigation whether heterozygote-specific structure in VariantFormer predictions reflects biological regulation.

We further assessed sensitivity to the size and source of the distillation genotype panel. Increasing the number of distillation-training individuals from 5 to 402 markedly improves distillation fidelity and increases the number of genes with well-fitted models, with improvements plateauing approximately at 200 individuals (**Appendix Figure 9c**). Using a size-matched panel of OneK1K genotypes for distillation (from the same individuals used in ATTICUS(+V) training) produce broadly similar fidelity distributions; ATTICUS distillation therefore does not depend on paired genotypes with its singe cell gene expression training data (**Appendix Figure 9d**).

### 3.5 Summary-TWAS on systemic lupus erythematosus and Alzheimer’s disease

We perform summary statistics TWAS (S-TWAS [15, 50]) on two disease traits, systemic lupus erythematosus (SLE) [56] and Alzheimer’s disease (AD), using ATTICUS(+V) distilled predictors and weights from models trained on OneK1K (containing blood cell-types) and ROSMAP (containing brain cell-types) respectively. For locus-level analyses, we used 1, 702 approximately independent linkage disequilibrium (LD) blocks [57]. We compared ATTICUS(+V) with l-ctPred in scPrediXcan, which we implement with the same GWAS summary statistics, in terms of the following metrics: enrichment for GWAS-implicated disease loci, recovery of previously identified TWAS-associated genes and LD blocks, cell-type specificity of association signals, and concordance of inferred effect directions (**Appendix Note F**).

For S-TWAS on SLE and AD using models trained on OneK1K and ROSMAP respectively, probabilistic, heterozygote-aware (HET) distillation of ATTICUS(+V) identifies more Bonferroni-significant genes across more distinct LD blocks (SLE: 168 significant genes across 24 LD blocks per cell-type; AD: 156 genes across 57 blocks, **Figure 3**) than either the deterministic HET distillation of ATTICUS(+V) (SLE: 115 genes across 20 blocks, AD: 100 genes across 48 blocks, **Appendix Figure 10**) or scPrediXcan (SLE: 130 genes across 18 blocks, AD: 108 genes across 47 blocks, **Appendix Figure 11**). We additionally perform cross-tissue sensitivity analyses using the ROSMAP-trained models for SLE and OneK1K-trained models for AD (**Appendix Figures 12, 13, 14**); crucially, these are not negative controls, since crosstissue disease-associated signals remain biologically plausible given e.g. peripheral T-cell involvement in AD [58] and microglia-mediated neuronal damage in experimental lupus [59]. Nonetheless, we expect more hits in blood cells for SLE and brain cells for AD, and we find that our probabilistic distillation method is the only one that shows this expected trend in both diseases (**Appendix Table 3**). Finally, we compare ATTICUS(+V) HET S-TWAS results with the dosage-only distillation (ADD) S-TWAS results, and find that they are comparable if not better in the former (**Figure 3a**, **Appendix Figures 15, 16**). From here on all ATTICUS(+V) S-TWAS results are from HET distillation.

We first assess whether the S-TWAS associations we have found include genes implicated by previous GWAS (**Figure 3a**; **Appendix Note F.1.2**). Across all 29 OneK1K cell-types tested for SLE and 31 brain cell-types tested for AD, S-TWAS using probabilistic ATTICUS(+V) distillation gives the highest mean GWAS gene set recovery: 12.6/117 tested genes for SLE (10.7%) and 28.7/179 for AD (16.0%), compared with 6.4 (5.5%) and 23.2 (13.0%), respectively, for scPrediXcan. This fraction is higher in genes where all distilled student models in probabilistic ATTICUS(+V) S-TWAS give Bonferroni-significant associations, suggesting that model agreement helps prioritise more robust associations (**Appendix Figure 17**). We then perform the same analysis using genes previously implicated by single cell TWAS [49] (**Appendix Note F, Appendix Table 4**); overall, probabilistic ATTICUS(+V) S-TWAS recovers a larger proportion of these associations than deterministic ATTICUS(+V) and scPrediXcan (**Appendix Figure 18**).

We further examine genomic inflation *λ*_GC_ in S-TWAS statistics separately for GWAS-implicated genes and all other non-implicated genes (**Figure 3a**): we expect this to be low and near 1 for non-implicated genes, and high for GWAS-implicated genes, assuming previous GWAS have identified most but not all genes relevant for each disease. As expected, probabilistic ATTICUS(+V) S-TWAS gives high mean *λ*_GC_ of 9.54 and 11.40 for GWAS-implicated genes in SLE and AD respectively, while giving low mean *λ*_GC_ of 1.34 and 1.23 for non-implicated genes in SLE and AD (**Figure 3a**). In comparison, both scPrediXcan and deterministic ATTICUS(+V) S-TWAS show lower signal for GWAS-implicated genes, and greater background inflation, with mean *λ*_GC_ values around 1.5 among genes outside the GWAS-implicated sets. Interestingly, we observe elevated inflation in the originally published scPrediXcan, consistent with previous reports of inflation in TWAS [60, 42, 61]: our re-analysis of the reported association *p*-values yields a mean *λ*_GC_ of 1.421 across 29 SLE cell-types and 1.628 across 11 T2D cell-types (**Appendix Table 5**). Altogether, these results show that our probabilistic ATTICUS(+V) model improves calibration and disease-specificity of S-TWAS signal through preferentially down-weighing associations driven by model-specific prediction noise. Disease-relevant signals more robust to variation across teacher models are preferentially retained in distillation and S-TWAS.

### 3.6 S-TWAS associations preserve cell-type specificity and effect direction

We next ask whether the improved prediction of cell-type-specific expression in ATTICUS(+V) (**Figure 2b**; **Appendix Figure 3b**) translates into better cell-type resolution in S-TWAS. We find that both deterministic and probabilistic ATTICUS(+V) S-TWAS identify more genes significant in exactly one of the 12 aggregated major cell-type groups than scPrediXcan. These genes also represented a larger proportion of all significant genes (**Appendix Table 6, Appendix Figure 19b**). Probabilistic ATTICUS(+V) S-TWAS identifies the most group-specific significant genes in both diseases, consistent with uncertainty-aware training and distillation contributing to improved cell-type-specific prediction and S-TWAS.

We then assess whether these group-specific S-TWAS hits are biologically plausible. For SLE (excluding MHC), probabilistic ATTICUS(+V) S-TWAS identifies the most group-specific hits in NK cells, closely followed by T and B cells (**Figure 3d**,**e**), consistent with previous reports implicating NK cells [62, 63] and T-cells [64] for SLE. For AD, probabilistic ATTICUS(+V) S-TWAS identifies the most specific hits in microglia (**Figure 3b**,**c**), consistent with AD genetic risk acting through microglial regulatory elements [65]. Neither cell-types are found to give the highest number of S-TWAS signal in scPrediX-can for their respective diseases (**Appendix Figure 11**). In cross-tissue analyses, we find the largest numbers of group-specific AD hits in blood T cells and platelets, consistent with reported T-cell and platelet involvement [58, 66]. For SLE, the choroid and ependymal group have the most group-specific hits, consistent with evidence implicating the choroid plexus as a site of lymphocyte accumulation and trafficking at the blood-CSF interface in neuropsychiatric lupus [67]. In comparison, scPrediXcan identifies only two platelet-specific AD hits and no choroid/ependymal-specific SLE hits (**Appendix Figures 12, 13, 14**). These results demonstrate that probabilistic ATTICUS(+V) S-TWAS has greater cell-type and disease specificity than previous approaches.

Finally, we assess effect-direction concordance of probabilistic ATTICUS(+V) S-TWAS findings against a curated set of gene–cell-type associations for SLE [68] and AD [69] whose effects on the diseases are identified through Mendelian Randomization (MR, **Appendix Note F.1.3**). Both deterministic and probabilistic ATTICUS(+V) S-TWAS show higher effect direction concordance than scPrediXcan in both SLE and AD gene sets, and the gain is particularly pronounced in AD (**Figure 3a**, two-sided binomial tests against 50% concordance deterministic: *p* = 0.0043; probabilistic: *p* = 0.0015). This is consistent with our finding that ATTICUS(+V) shows improved personalized gene expression prediction sign accuracy for some genes and cell-types, where effects are shared with bulk tissues (**Appendix Figures 5, 7, 8**).

## 4 Discussion

In this paper, we propose ATTICUS, a deep MLP ensemble that transfers variant-aware representations learnt in bulk-tissue S2F models such as VariantFormer [36] to personalised, cell-type-specific gene expression prediction without requiring matched genotypes for the single-cell training data. ATTICUS improves cell-type-specific prediction accuracy relative to scPrediXcan [21] and achieves high sign accuracy in personalised predictions for gene–cell-type pairs whose regulatory effects agree with those in the corresponding bulk tissue. Heterozygosity-aware probabilistic distillation propagates predictive uncertainty into downstream summary-TWAS, yielding more significant associations with reduced background genomic inflation. Though subject to limitations **Appendix Note G**, ATTICUS enables the use of any ungenotyped single-cell atlases with substantial donor numbers sizes [5, 6, 70] for cell-type specific genetic association analysis.

Most notably, we find that using variant-aware S2F backbones trained on bulk expression, including VariantFormer [36] and Variformer [34], substantially improves the directional consistency of personalised cell-type-level pseudobulk expression predictions in ATTICUS compared with reference-trained Enformer [16]. Predicted single variant effects from both reference-based and variant-aware S2Fs have been shown to have strong correlations between distilled variant effects and eQTL effects [16, 71, 34, 36], yet reference-trained models often show limited sign accuracy in personalised expression prediction [26, 24, 25, 72]. This discrepancy is consistent with a recent report showing that underestimation of causal variant effects relative to surrounding background predictions can leave personalised predictions dominated by noisy contributions [29]. It seems possible that inter-individual variations in variant effects are too small to be learnt using across-genes sequence contexts from a reference genome. It remains to be investigated whether improved learning of regulatory grammar and distal regulatory interactions can overcome this limitation, or whether explicit training on population-level genetic variation is required.

Further, we find that improvements in sign accuracy in ATTICUS are largely restricted to gene–cell-type pairs whose regulatory effects agreed with bulk-derived predictions (**Appendix Figure 7**), and where genes have high cell-type-shared cis-heritability (**Appendix Figure 8**). This suggests that transfer is most effective for cell-type shared regulatory effects learnt by S2Fs pretrained with bulk-tissue gene expressions, and least effective for cell-type specific ones. This limitation matters because the latter is where constraint has been found to be lower [3] and complex-trait heritability has been found to be enriched [4]. These cell-type specific effects are masked in bulk expression measurements and eQTL studies [2], leading to high TWAS false sign rates [73], and our findings show that the same problem plagues bulk-trained S2F backbones. Therefore, our findings suggest the need for cell-type resolved S2F model training for it to capture cell-type specific effects most likely to be relevant to complex traits and diseases.

Last but not least, a central contribution of ATTICUS is the propagation of predictive uncertainty from a deep ensemble [48] through probabilistic distillation into S-TWAS [50, 53], resulting in more TWAS hits that are supported by GWAS [74, 75] and TWAS [49] findings (**Figure 3a, Appendix Figure 18**). These gains were accompanied by clearer separation between established disease signals and background associations (**Figure 3a**), consistent with previously reported benefits of accounting for predictive uncertainty in TWAS [44, 43, 45]. However, our uncertainty estimates are conditional on a single frozen S2F backbone and do not represent uncertainty in its learnt representations. It has recently been shown that uncertainty in direction concordance (and subsequently also in TWAS effect direction) can be captured via cross-validation replicates of the same S2F model [29], which is something that cannot be captured by a single S2F backbone in ATTICUS. As such, propagating variation across independently trained backbones may improve the characterisation of directional uncertainty in future versions of ATTICUS.

Altogether, ATTICUS demonstrates that single-cell datasets without matched genotypes can be used to derive personalised expression predictors by transferring regulatory information from bulk-trained S2F models. Coupling these predictors with uncertainty-aware distillation and association testing provides a practical route to cell-type-specific TWAS and other gene-level analyses across the expanding landscape of single-cell atlases [5, 6, 70], including tissues and cell populations for which sufficiently large (e.g. biobank scale) genotyped expression cohorts are and will likely remain unavailable.

## 5 Code Availability

ATTICUS is available at https://github.com/Padraig20/atticus.

## 6 Author Contributions

P.S. implemented the methods. P.S. and Y.D. analyzed the data. P.S., Y.D and N.C. interpreted the results. P.S. and Y.D. and N.C. wrote the manuscript. N.C. conceived the study and supervised the work.

## 7 Acknowledgements

NC is supported by the Helmut Horten Foundation Young Investigator Grant 2025-YIG-013 and the Swiss National Science Foundation Project Grant CR00-5-242822.

The results published here are in part based on data obtained from The AD Knowledge Portal: genomic and scRNAseq data from brain tissues used in this study were generated from postmortem brain tissue provided by the Religious Orders Study and Rush Memory and Aging Project (ROSMAP) cohort at Rush Alzheimer’s Disease Center, Rush University Medical Center, Chicago. Data collection was supported through funding by NIA grants P30AG10161 (ROS), R01AG15819 (ROSMAP; genomics and RNAseq), R01AG17917 (MAP), R01AG30146, R01AG36042 (5hC methylation, ATACseq), RC2AG036547 (H3K9Ac), R01AG36836 (RNAseq), R01AG48015 (monocyte RNAseq) RF1AG57473 (single nucleus RNAseq), U01AG32984 (genomic and whole exome sequencing), U01AG46152 (ROSMAP AMP-AD, targeted proteomics), U01AG46161(TMT proteomics), U01AG61356 (whole genome sequencing, targeted proteomics, ROSMAP AMP-AD), P30AG072975, the Illinois Department of Public Health (ROSMAP), and the Translational Genomics Research Institute (genomic). Specifically the scRNAseq data was funded by NIH grants U01AG061356 (De Jager/Bennett), RF1AG057473 (De Jager/Bennett), and U01AG046152 (De Jager/Bennett) as part of the AMP-AD consortium, as well as NIH grants R01AG066831 (Menon) and U01AG072572 (De Jager/St George-Hyslop). The authors thank all members of the ROSMAP analytic team for making this data available, and all participants who contributed to the ROSMAP cohorts for making this work possible.

The results published here are in part based on data obtained from the 1000 Genomes and OneK1K Consortium, which are both released openly and publicly for the benefit of the wider biomedical community. The authors thank all members of the 1000 Genomes and OneK1K Consortium for making this data available, and all participants who contributed to the 1000 Genomes and OneK1K for making this work possible.

The authors thank Katherine Pollard and her team for providing the Variformer model weights for GTEx tissues upon request and demonstration of controlled access to GTEx RNAseq data.

The authors thank Valentina Boeva, Lovro Rabuzin, Luca Pinello, Andy Dahl, Laramie Duncan and Jonathan Pritchard for helpful discussions and feedback.

## 8 Use of AI

We used large language models and AI coding agents for drafting, language editing, and assistance with software development and data analysis workflows. All AI-generated code and analyses were reviewed and validated by the authors. The final responsibility for the manuscript, including its scientific contributions, interpretation of results and presentation, remains with the authors.

## 9 Conflicts of interest

The authors declare no conflicts of interest.

## Notes

### Competing Interest Statement

The authors have declared no competing interest.

