## Supplementary Materials for "Uncertainty-Aware Personalised Gene Expression Prediction Improves Cell-Type-Specific Association Testing"

#### Appendix contents

**A Ethical Approval** **S2**

**B Implementation Details** **S2**

B.1 Paired genetic and single-cell transcriptomic data . . . . . S2
B.2 Model training . . . . . S2
B.3 Model distillation . . . . . S3
B.4 GWAS summary statistics for S-TWAS . . . . . S4
B.5 Summary TWAS . . . . . S4

**C Gaussian Likelihood Analysis of Reference-Based Training** **S6**

**D Instability of the $m_1$ Metric at Small Non-Null Fractions** **S7**

**E What affects cross-individual prediction** **S8**

E.1 Effect of variant-aware, tissue-conditioned training . . . . . S8
E.2 Effect of personal-genome training vs model-specific architecture . . . . . S8
E.3 Relationship between prediction uncertainty and prediction accuracy . . . . . S8

**F Summary TWAS evaluations** **S9**

F.1 Recovery of GWAS-implicated genes and comparison with scPrediXcan . . . . . S9
F.2 Gene and cell-type concordance with published scTWAS findings . . . . . S10

**G Limitations** **S11**

G.1 Unseen genes . . . . . S11
G.2 Portability . . . . . S11
G.3 Evaluation standards . . . . . S11
G.4 Spurious or undiscovered S-TWAS genes . . . . . S11
G.5 Cell-type specificity . . . . . S11
G.6 Uncertainty about direction . . . . . S12

**H Appendix Tables** **S13**

**I Appendix Figures** **S20**

### A Ethical Approval

The ROSMAP [1, 2, 3] data is obtained through controlled access approval from the AD Knowledge Portal (synID: syn3219045).

### B Implementation Details

#### B.1 Paired genetic and single-cell transcriptomic data

We used two scRNA-seq datasets with genotype data for our model training and validation. First, we obtained pre-processed OneK1K single-cell RNA-seq (scRNA-seq, 10,385–16,193 genes per cell-type, 29 cell-types, in mean raw UMI counts per cell) and genotype data (8,439,610 SNPs with MAF $> 1\%$) for 982 individuals as previously described [4] from the official CELLxGENE repository[1]. We used a total of 17,967 protein-coding genes across 29 cell-types in our experiments, which is the union of genes present in at least one cell-type. Unlike the data used in the original scPrediXcan study, which were specifically pre-processed by the Powell laboratory, we did not apply additional pre-processing beyond aggregation to pseudo-bulk profiles. Differences in pre-processing may therefore contribute to the difference between the prediction performance observed in our experiments and that reported for scPrediXcan. Second, we obtained pre-processed single-nucleus RNA-seq (snRNA-seq, 12–16,402 genes per cell-type, 31 Allen MapMyCells superclusters in mean raw UMI counts per cell) and genotype data (10,442,858 SNPs with MAF $> 1\%$) for 396 individuals labelled as controls (non-Alzheimer's Disease patients) in the ROSMAP dataset. The variant positions in the genotype data, originally in GRCh37/hg19, were mapped to GRCh38/hg38 using the UCSC `liftOver` tool. We used a total of 15,915 protein-coding genes (union of genes present in at least one supercluster) across 31 cell-type superclusters, 21 of which are neuronal.

We used single cell gene expression data for 72% of genes in 80% of individuals in each dataset for reference training in ATTICUS, and assess the model prediction accuracy in three ways as shown in **Figure 2**: a) Pearson's correlation of prediction and observed single cell gene expression across genes within each cell-type in the held out 13% of test-genes, averaged across all individuals used in training for plotting in **Figure 2b**; b) Pearson's correlation of prediction and observed single cell gene expression across cell-types for each gene in the held out 13% of test genes, averaged across all individuals used in training for plotting in **Figure 2b**; and c) Pearson's correlation of prediction and observed single cell gene expression across individuals for each gene in each cell-type in the held out 20% of test individuals, averaged across all genes per cell-type in **Figure 2d** and **Appendix Figure 3a,b**.

#### B.2 Model training

##### B.2.1 ATTICUS(+V)

We used the sequence-to-function (S2F) model VariantFormer [5] to encode the cis-regulatory sequence context surrounding each gene. This S2F backbone is frozen, meaning its parameters are not updated during training of the downstream multi-layer perceptron (MLP). For each gene, we obtain the human genome reference GRCh38 sequences around it, including all candidate cis-regulatory elements (cCREs) within 1Mb of the gene and a transcription window extending from 1kb upstream of its TSS to 300kb downstream (or the gene end). VariantFormer integrates these sequences with tissue context to produce a 1,536-dimensional tissue-conditioned gene representation. Whereas Enformer uses a single DNA sequence and is not diploid-aware, VariantFormer directly represents personalised diploid genomes and incorporates heterozygous and homozygous genetic variants into its sequence representation. We extracted embeddings from the protein-coding VariantFormer checkpoint (`variantformer-pcg`), which was trained to predict the expression of 18,439 protein-coding genes across diverse tissues and cell lines. Embeddings were conditioned on the `blood` tissue token for blood cell-types in the OneK1K dataset, and on the `brain - frontal cortex (ba9)` tissue token for all ROSMAP DLPFC cell-types. We used genes in chromosomes 1, 2, 3, 4, 6, 8, 9, 10, 13, 15, 16, 17, 18, 19, 21, 22, X and Y for training; 7, 11 and 14 for validation; 5, 12, 20 for testing. This is consistent with the split used by the method we benchmark ourselves against, scPrediXcan [6].

We fed the tissue-conditioned gene representations into an MLP ensemble with a total number of $K = 5$ members. Each member has the same layout, and they differ only in weight initialisation. The number

[1] `https://cellxgene.cziscience.com/collections/dde06e0f-ab3b-46be-96a2-a8082383c4a1`

of hidden layers can be freely specified, and the size of the separate layers corresponds to a linear interpolation between the input and output layer, with optional layer normalisation over a mini-batch before the first hidden layer. Each hidden layer is followed by a ReLU activation function. To ensure positive predictions for both mean and variance, we added Softplus activation after the output layer. The hidden layers of our MLP are non-identical and their parameters are not shared within or across ensemble members. During training, we added a learning rate scheduler with linear warmup and cosine decay, as well as an early stopping mechanism (patience = 5) by monitoring validation loss. The deep ensembles were saturated after roughly 20 epochs.

###### B.2.2 ATTICUS(+E)

To benchmark against our full model ATTICUS(+V), we also assessed performance when the S2F backbone is switched to Enformer [7], the S2F used in scPrediXcan. Here we followed exactly the same procedure as ctPred in scPrediXcan [6]: we provide a 196,608-bp sequence centred on the TSS, from which Enformer produces an $896 \times 5{,}313$ representation. If the transcription start site (TSS) cannot be centred within the input sequence without padding, we shift the sequence window to minimise padding [8]. We average the four central bins (447–450) to obtain a 5,313-dimensional gene representation. All model training processes are the same as described above.

###### B.2.3 ctPred and ctPred* benchmarks

The ATTICUS architecture is highly similar to the ctPred model as stated in both the scPrediXcan [6] paper and its official implementation: an initial linear layer that maps the Enformer embedding to a hidden layer of 64 dimensions, followed by ReLU, dropout, and three further identical hidden layers with parameter-sharing to map to the final predictions. This enables us to compare our model to ctPred in downstream analyses. As compared to our deep ensembles which saturated after 20 epochs, ctPred models required at most 100 epochs, consistent with reports from the scPrediXcan manuscript.

###### B.2.4 Prediction target and loss functions

One important difference between ATTICUS and ctPred is that ctPred aims to predict the percentile of a gene's expression when ranked against all other genes in each cell-type. Specifically, the ctPred model requires a specific transformation of the cell-type-specific gene-by-individual pseudo-bulk matrix: genes are ranked by their average pseudo-bulk expression across individuals, and the ranks are then converted into percentiles ranging from 0 to 1. The ctPred model was trained using a loss function for Mean Squared Error (MSE) alone.

Each member of our deep MLP ensemble in ATTICUS(+V), on the other hand, predicts log-transformed pseudo-bulk gene expression directly. This is a fundamentally more difficult prediction task, as it requires recovering both the relative ordering of genes and the quantitative differences in their expression levels, whereas percentile-based targets retain only relative ranks. To account for this increased difficulty, we used a different loss function inspired by the UNICORN model [9], which consists of a learnt weighted mixture of Pearson correlation (PC), Cosine Similarity (CS), Poisson Negative Log Likelihood (PNLL) and Gaussian Negative Log Likelihood (GNLL). GNLL serves as the sole estimator of predictive variance, and is otherwise equivalent to MSE.

##### B.3 Model distillation

###### B.3.1 Distillation genetic data

For distillation of the MLP deep ensemble (the teacher model) into a linear student model whose effects can be directly used in summary TWAS (S-TWAS), we used genotype data from the 1000 Genomes Project (1000G) Phase 3 data [10], obtained from the genotype resources released with scPrediXcan[2]. We restricted the data to 6,000,825 SNPs with a minor allele frequency (MAF) of $\geq 5\%$ in 503 individuals of European ancestry as stated on the 1000G metadata file; scPrediXcan used a total of 462 European individuals from the 1000G cohort.

[2] https://zenodo.org/records/16538482

##### B.3.2 Probabilistic distillation

We first generated predictions for the 503 individuals we used from the 1000G cohort by using the MLP deep ensemble. For each of the $K = 5$ ensemble members, we saved its normalised gene expression predictions (inverse normal transform across all individuals) separately, including predictive mean and variance. We then distilled each member separately, using an 80:20 individual split, where the 80% training fold is used for bootstrapping. For each of the $K = 5$ ensemble members, we bootstrapped $B = 5$ elastic nets, where each is a standard with-replacement draw of size corresponding to the 80% training individuals used for distillation. Since VariantFormer, which we used in ATTICUS(+V), has a maximum sequence window of 1Mb, we restricted the student model to SNPs within the same genomic window. Including variants outside the teacher's receptive field would introduce predictors that could not have contributed to the teacher's expression estimate and would therefore be inconsistent with the distillation setup. We arrived at a total of $B \times K = 25$ linear models, which we used to derive predictions in individuals in the 20% test fold and then average across them. Note that this is equivalent to deriving a single prediction using a model with average weights from all $B \times K = 25$ linear models. We assessed whether the number of individuals' genotypes we used in distillation affected distillation efficacy, increasing the number of training individuals in distillation from 5 to 402 (the full 80%).

##### B.3.3 Heterozygosity-aware distillation and additive projection

Unlike Enformer, which is used by scPrediXcan and processes single DNA sequences, VariantFormer explicitly represents diploid genotypes. To accommodate heterozygote-specific effects in its predictors, we fitted elastic-net students using both SNP dosages and heterozygosity indicators. For each gene–cell-type pair, the student prediction was

$$\hat{y} = \sum_j \beta_j G_j + \sum_j \delta_j \mathbb{1}(G_j = 1), \tag{1}$$

where $G_j \in \{0, 1, 2\}$ is the counted allele dosage at SNP $j$, and $\delta_j$ allows the heterozygous contribution to deviate from the midpoint of the two homozygous contributions. We estimated both coefficient sets jointly using the uncertainty-weighted elastic-net objective.

To obtain additive predictors compatible with S-PrediXcan, we subsequently converted the fitted coefficients into population-average allele-substitution effects. Under Hardy-Weinberg equilibrium, the effective additive weight for SNP $j$ is

$$\beta_j^{\text{eff}} = \beta_j + (1 - 2p_j)\delta_j, \tag{2}$$

where $p_j$ is the frequency of the allele counted by $G_j$, estimated from the genotype reference panel. This projection incorporates the contribution of heterozygote-specific effects to the average additive effect, and makes the predictor compatible with downstream association testing [11]. Hence, we applied it separately to each of the $K \times B$ students. The resulting additive weights, together with their SNP covariance matrices, were used for S-PrediXcan association testing. No heterozygosity-specific GWAS summary statistics are required in S-TWAS using these distilled weights.

#### B.4 GWAS summary statistics for S-TWAS

We used summary statistics for GWAS on SLE performed on 14,267 individuals (5,201 SLE cases, 9,066 controls, GWAS Catalog entry `GCST003156`) of European ancestry [12] and GWAS on AD performed on 1,119,443 individuals (75,638 AD cases, 1,043,805 controls, GWAS Catalog entry `GCST90704647`) of European ancestry [13] to perform S-TWAS. As our distilled model only contains SNPs with minor allele frequency (MAF) of $\geq 5\%$ in 1000G Phase 3 Europeans, and would only use these SNPs for S-TWAS, we do not perform additional filtering on the GWAS summary statistics.

#### B.5 Summary TWAS

We used the S-PrediXcan framework [14] to perform S-TWAS. Each of the distilled $B \times K = 25$ linear models, given same GWAS data, have been used with S-PrediXcan to acquire $B \times K = 25$ different S-TWAS outcomes. $p$-values were combined using aggregated Cauchy association test (ACAT) [15, 16], which works well under arbitrary correlation structures, while the other statistics were simply averaged.

S-PrediXcan requires estimates of SNP variances, which we estimated using the same genotype data used for distillation. A SNP covariance matrix is provided with each model we trained. For locus-level analyses, we used the approximately independent linkage disequilibrium (LD) blocks [17]; of the 1,703 LD blocks originally provided in hg19 coordinates, 1,702 were successfully mapped to hg38 using the UCSC `liftOver` tool.

#### C Gaussian Likelihood Analysis of Reference-Based Training

During reference-based training, all individuals share the same sequence representation for a given gene $g$, while their observed cell-type-specific pseudo-bulk expression values $\{y_{igc}\}_{i=1}^{N}$ may differ. Thus, for a fixed gene $g$ and cell-type $c$, the probabilistic predictor receives the same input $\mathbf{h}_g^{\text{ref}}$ for all individuals from the S2F backbone, but is trained against $N$ distinct expression measurements $\{y_{igc}\}_{i=1}^{N}$.

Consider the Gaussian likelihood used by the GNLL component of our training objective. Conditional on the reference embedding, assume

$$y_{igc} \mid \mathbf{h}_g^{\text{ref}} \sim \mathcal{N}(\mu_{gc}, \sigma_{gc}^2), \qquad i = 1, \dots, N, \tag{3}$$

with observations conditionally independent across individuals. Because the input is identical for all individuals, the model assigns the same $\mu_{gc}$ and $\sigma_{gc}^2$ to each observation. The joint log-likelihood for a fixed $(g, c)$ is

$$\mathcal{L}(\mu_{gc}, \sigma_{gc}^2) = \sum_{i=1}^{N} \log \mathcal{N}\left(y_{igc}; \mu_{gc}, \sigma_{gc}^2\right) \tag{4}$$

$$= -\frac{N}{2} \log(2\pi\sigma_{gc}^2) - \frac{1}{2\sigma_{gc}^2} \sum_{i=1}^{N} (y_{igc} - \mu_{gc})^2. \tag{5}$$

Differentiating with respect to $\mu_{gc}$ gives

$$\frac{\partial \mathcal{L}}{\partial \mu_{gc}} = \frac{1}{\sigma_{gc}^2} \sum_{i=1}^{N} (y_{igc} - \mu_{gc}). \tag{6}$$

Setting the derivative to zero yields

$$\hat{\mu}_{gc} = \frac{1}{N} \sum_{i=1}^{N} y_{igc} = \bar{y}_{gc}. \tag{7}$$

Similarly, differentiating with respect to $\sigma_{gc}^2$ and substituting $\hat{\mu}_{gc}$ gives

$$\hat{\sigma}_{gc}^2 = \frac{1}{N} \sum_{i=1}^{N} \left(y_{igc} - \bar{y}_{gc}\right)^2. \tag{8}$$

Thus, under Gaussian likelihood, repeatedly pairing the same reference embedding with individual-specific expression measurements yields two different population-level quantities: the mean expression of gene $g$ in cell-type $c$ and the variation of expression across individuals. Collapsing the observations to $\bar{y}_{gc}$ before training would retain information about the mean identically but discard the variation required to estimate $\sigma_{gc}^2$. A simplified schematic of training the S2F-based expression predictor is provided in **Appendix Figure 1**.

#### D Instability of the $m_1$ Metric at Small Non-Null Fractions

We saw, empirically, that the $m_1$-metric proposed by scPrediXcan [6] is incredibly unstable across seeds despite predictions barely changing (**Appendix Figure 2**). We show that this happens when the signal (of the alternative hypothesis $\pi_1$) is weak compared to the estimated null hypothesis $\pi_0$. The estimated number of true positive (non-null) genes is defined as $m_1 = m(1 - \pi_0)$, with $m$ being the total number of genes. Let $\delta$ be an infinitesimal perturbation in $\pi_0$.

$$
\begin{aligned}
m_1' &= m(1 - (\pi_0 + \delta)) && (9)\\
&= m(1 - \pi_0 - \delta) && (10)\\
&= m(1 - \pi_0) - m\delta && (11)\\
&= m_1 - m\delta && (12)
\end{aligned}
$$

Then, the difference between $m_1$ and the perturbed $m_1'$ can be expressed as the relative error:

$$
\begin{aligned}
&\Delta m_1 = m_1' - m_1 = -m\delta && (13)\\
\Leftrightarrow\quad &\frac{|\Delta m_1|}{m_1} = \frac{|\delta|}{1 - \pi_0} && (14)
\end{aligned}
$$

Naturally, this diverges as $\pi_0 \to 1$. In our setting, most genes have weak correlations, so their $p$-value distribution is close to uniform: only a small fraction of genes deviate from the null, such that $\pi_1 = 1 - \pi_0 \ll 1$. Small seed-induced changes in the predictions can therefore cause small changes in the empirical $p$-value distribution and therefore in the estimated $\pi_0$.

### E What affects cross-individual prediction

#### E.1 Effect of variant-aware, tissue-conditioned training

Our main experiments showed that variant-aware, tissue-conditioned VariantFormer representations in ATTICUS(+V) improve both reference expression prediction and, more substantially, the sign consistency of personalised predictions compared with reference-based Enformer in ATTICUS(+E). However, VariantFormer differs from Enformer in several ways, making it difficult to determine which modelling choices actually drive these improvements without detailed ablation studies. We therefore hypothesize how its learnt representations may be an improvement over those in Enformer for the task of personalized single cell gene expression prediction.

VariantFormer is a hierarchical transformer trained on 21,004 paired genome–bulk-transcriptome samples from 2,330 donors. In addition to explicitly representing diploid genetic variation as input, it incorporates a gene-specific transcriptional sequence window together with candidate cis-regulatory elements within 1 Mb and conditions its predictions on tissue identity, rather than a fixed cis-window of regulatory context used in Enformer. Thus, its improvements over Enformer could arise from population-scale genome–transcriptome training, tissue conditioning, its representation of diploid variation, its more comprehensive sequence regulatory context that encompasses distal enhancers not utilised in Enformer, or other architectural differences.

#### E.2 Effect of personal-genome training vs model-specific architecture

We additionally considered Variformer [18], which is architecturally a lot closer to Enformer. Indeed, Variformer keeps Enformer's architecture, dense sequence context and one-hot haplotype encoding, but is fine-tuned on personal genomes paired with whole-blood expression for about 300 genes. We therefore compare using Variformer to both Enformer and VariantFormer as the S2F backbone in ATTICUS: the Enformer–Variformer comparison is a near-direct test of the effect of personal-genome fine-tuning; VariantFormer-Variformer comparison instead shows what personal-genome training achieves at scale, across all protein-coding genes and 54 tissues, though it cannot separate the contribution of the training data from VariantFormer's other differences in architecture, sparse long-range regulatory context, diploid encoding and tissue conditioning.

We restricted this analysis to the 300 genes used for Variformer Whole Blood fine-tuning and trained otherwise identical downstream predictors from Enformer, Variformer, and VariantFormer representations using the same protocol as in our main experiments. We will refer to our Variformer-based method as ATTICUS(+VE). Both ATTICUS(+VE) and ATTICUS(+V) improve prediction of held-out OneK1K genes relative to ATTICUS(+E) (**Appendix Figure 6a**). More strikingly, while the magnitude of across-donor correlations improves only modestly for both population-trained models (**Appendix Figure 6b**), they substantially improve the sign consistency of predicted expression differences across individuals, whereas ATTICUS(+E) remains close to random (**Appendix Figure 6c**). Together, these results suggest that training on paired genetic variation and bulk expression across individuals is an important factor underlying the improved personalised predictions of ATTICUS(+V), while its additional architectural and large-scale pre-training differences may provide further gains.

#### E.3 Relationship between prediction uncertainty and prediction accuracy

High *epistemic* uncertainty estimated in the teacher model in ATTICUS (**Methods**) identifies genes whose overall expression is difficult to predict by our models, while high *aleatoric* uncertainty identifies genes whose variance is high in the training population. However, we found that both uncertainty estimates only partially identify which genes will exhibit less reliable cross-individual predictive performance. Nonetheless, we can reliably decide which individuals are harder to predict: mean within-gene, cross-individual Spearman correlations between aleatoric uncertainty and squared prediction error ranged from 0.35 to 0.76 across OneK1K cell-types and from 0.00 to 0.41 in ROSMAP cell-types; associations with epistemic uncertainty were slightly weaker (**Appendix Figure 4**).

### F Summary TWAS evaluations

#### F.1 Recovery of GWAS-implicated genes and comparison with scPrediXcan

##### F.1.1 LD blocks rather than individual genes.

Additional S-TWAS gene discoveries in ATTICUS(+V) S-TWAS as compared to scPrediXcan may be attributed to more sequence regulatory effects learnt through the more varied sequence context (gene-specific CREs, TSS) in the S2F we used in ATTICUS(+V), VariantFormer, rather than a consistent cis-window used in the scPrediXcan's S2F, Enformer. If so, this may result in ATTICUS(+V) S-TWAS identifying more significant genes residing in distinct loci from those in scPrediXcan. It may also be attributable to effects learnt from the same sequence context, through an improvement in transfer of effects learnt from bulk RNA-seq training of either S2F to cell-type level predictions enabled by our uncertainty-aware deep ensemble training architecture and uncertainty-preserving distillation process prior to S-TWAS. If so, this may result in ATTICUS(+V) S-TWAS identifying more significant genes in the loci that also contain S-TWAS hits in scPrediXcan.

To assess whether additional S-TWAS hits arise within the same loci or extend to other loci, we grouped Bonferroni-significant gene–cell hits by previously defined LD blocks. Probabilistically distilled ATTICUS(+V) HET identified more hits than scPrediXcan in SLE (168.4 versus 130.1 per cell-type; paired t-test $p = 9.9\times10^{-20}$) and AD (155.6 versus 108.3; $p = 1.8\times10^{-21}$). Its hits also spanned more LD blocks per cell-type: 24.1 versus 18.4 in SLE ($p = 1.3 \times 10^{-10}$) and 57.1 versus 46.9 in AD ($p = 1.3 \times 10^{-13}$). Most ATTICUS-specific hits fell in blocks that also contained a scPrediXcan hit (86% in SLE and 74% in AD), while the higher block counts show additional locus coverage. The high SLE overlap is influenced by the MHC: outside that region, 50% of ATTICUS-specific hits fell in scPrediXcan-positive blocks. Within the ATTICUS(+V) model family, probabilistic HET also exceeded deterministic HET in both hits and block coverage, consistent with a contribution from the uncertainty-aware training and distillation pipeline. This attests to our hypothesis that our uncertainty-aware deep ensemble training architecture and uncertainty-preserving distillation process in ATTICUS enables the identification of more hits given the same sequence context.

##### F.1.2 GWAS implicated gene identification

To evaluate whether our S-TWAS had performed well and identified genes previously found to be associated with the diseases, we derived "GWAS-implicated genes" using 121 genes reported as candidate genes at SLE-associated loci [19] and 188 genes nominated as putative effector genes at AD-associated loci [20], respectively.

To assess if we obtain enriched ATTICUS(+V) S-TWAS signal in GWAS implicated genes, we stratified all genes we test in S-TWAS for SLE and AD into a "GWAS-implicated" set (as described above) and an "all others" set containing all remaining genes. We examined TWAS $p$-value distributions from ATTICUS(+V) S-TWAS and results from scPrediXcan within each stratum and comparable cell population. We ran scPrediXcan ourselves rather than using their published results, so that we have a fair comparison using the same input GWAS for S-TWAS for both diseases. We calculated ATTICUS(+V) and scPrediXcan S-TWAS genomic control index $\lambda_{\text{GC}}$ for each set of genes for each cell-type tested for SLE and AD as the median one-degree-of-freedom chi-squared statistic corresponding to the association $p$-values divided by the null median of approximately 0.455. We then assessed whether association evidence was preferentially concentrated among GWAS-implicated genes, rather than showing a similar distributional shift among all remaining genes; we assume genes previously implicated by GWAS are more likely to be truly relevant to disease, while genes not previously implicated are less likely so. As such, we expect to see higher $\lambda_{\text{GC}}$ values among GWAS-implicated genes and lower $\lambda_{\text{GC}}$ values among other genes in a S-TWAS that captures disease-relevant signal.

##### F.1.3 Direction of effect concordance with previous findings

To assess the direction of S-TWAS associations, we assembled a reference for directions of gene effects on SLE and AD from published cell-type-specific genetic analyses. For SLE, we used single-cell cis-eQTL-based Mendelian randomisation (MR) and colocalisation results across 14 immune cell-types from [21]. This study combined OneK1K cis-eQTLs with FinnGen R12 SLE GWAS summary statistics (1,306 cases and 372,273 controls), whereas our S-TWAS used the SLE GWAS by Bentham et al. [19](GCST003156,

5,201 cases and 9,066 controls). We retained all 62 gene–cell-type associations reported as MR-significant in Supplementary Table S1 of [21], using the sign of the MR effect estimate as the reference direction. For AD, we used cell-type-specific Summary MR (SMR) and colocalisation results from [22], which integrated multiple brain eQTL resources (including ROSMAP) with five AD GWAS datasets, all of which share contributing cohorts with the GWAS we used in S-TWAS. Here, we also retained all gene–cell-type pairs, reported as passing SMR FDR $< 0.05$, $P_{\text{HEIDI}} > 0.05$, PP.H4 $> 0.75$, and PP.H4/PP.H3 $> 3$.

These selected gene–cell-type pairs were mapped to the corresponding gene–cell-type pairs predicted and distilled from OneK1K and ROSMAP models in ATTICUS(+V), which we used for S-TWAS on SLE and AD, respectively. Positive MR/SMR effects indicated increased disease risk with higher genetically predicted expression, and negative effects indicated reduced risk. We compared these reference signs with signed S-TWAS estimates among all pairs meeting the specified S-TWAS significance threshold. Where a reference cell class mapped to multiple finer clusters, concordance was interpreted at the broader cell-class level.

Notably, both references used eQTL data originating from the same gene expression datasets we use in our model training (OneK1K and ROSMAP) and the AD reference used GWAS that shared cohorts with the AD GWAS we use in our S-TWAS. We therefore interpret sign agreement as directional concordance rather than independent causal validation. Repeated genes across cell-types and results across overlapping GWAS were not treated as independent biological replications.

##### F.2 Gene and cell-type concordance with published scTWAS findings

We compared our SLE and AD TWAS results with published scTWAS gene–cell-type associations [23], using Supplementary Data 3 and 5, respectively. Both analyses used the same SLE GWAS summary statistics [12], whereas the AD GWAS differed between our study [13] and scTWAS [24]. We mapped 12 published SLE cell-types and six AD major cell-types to compatible populations in our analyses (**Appendix Table 4**), excluding SLE plasma cells because no corresponding population was available. Where a published population corresponded to multiple cell-types in our analyses, we combined their gene-level $p$-values using ACAT. Significance was assessed using Bonferroni correction across all genes tested within each matched population, consistent with the correction used by scTWAS. Of the 143 SLE and 81 AD published gene–cell-type associations retained after cell-type matching, 131 and 69, respectively, were evaluable.

Probabilistic ATTICUS recovered a larger proportion of these associations than deterministic HET and scPrediXcan (**Appendix Figure 18a**). For SLE, probabilistic ADD and HET recovered 84/131 (64.1%) and 83/131 (63.4%) associations, respectively, compared with 61/131 (46.6%) for deterministic HET and 65/131 (49.6%) for scPrediXcan. For AD, the corresponding recovery rates were 35/69 (50.7%), 29/69 (42.0%), 24/69 (34.8%), and 12/69 (17.4%). At the gene level, probabilistic ADD and HET recovered 29/34 (85.3%) and 30/34 (88.2%) evaluable SLE genes, respectively. Recovery was lower for AD, at 24/45 (53.3%) and 22/45 (48.9%). The difference between diseases should be interpreted in the context of their different GWAS inputs and cell-type mappings.

We also inspected significant genes absent from the supplied scTWAS hit lists across all 29 original SLE and 31 original AD cell-types (**Appendix Figure 18b**). Probabilistic ADD identified 308 such SLE genes and 423 AD genes, representing 90.9% and 94.6% of its distinct significant genes, respectively. Probabilistic HET identified 299 SLE genes (90.9%) and 410 AD genes (94.5%). Most of these genes were significant in at least two tested cell-types; the figure separately reports genes significant in exactly one cell-type. These findings show that ATTICUS recovers published associations while identifying a broader set of candidate genes. However, absence from the supplied scTWAS hit lists does not establish disease-gene novelty, and significance in only one tested cell-type does not establish biological cell-type specificity.

### G Limitations

Several limitations should be considered when interpreting our findings.

#### G.1 Unseen genes

Although we evaluated the transfer of learnt cell-type level gene representations to held-out genes in the ATTICUS training step, these genes were excluded only from cell-type-specific training and not from S2F [5] pretraining. Our results therefore demonstrate transfer to previously unseen cell-type-specific expression targets rather than generalization to genes entirely unseen by the S2F backbone.

#### G.2 Portability

Both our cell-type specific training and our genotype-based distillation and downstream S-TWAS analyses were restricted to European-ancestry data [4, 1, 2, 3, 10, 12, 13], leaving portability across ancestries untested. Differences in allele frequencies, linkage disequilibrium and genetic backgrounds may affect both the distilled prediction weights and downstream association results. The same is true for portability between cell-types of different developmental stages and disease contexts.

#### G.3 Evaluation standards

Our disease-specific evaluation standards [21, 22, 20, 19, 23] included genes implicated by previous GWAS or scTWAS analyses. Some contributing cohorts overlap those used in our S-TWAS analyses or model training, so concordance measures agreement with existing evidence rather than independent replication. In terms of recovery of previously identified GWAS hits, GWAS-implicated gene sets we used for this evaluation [19, 20] are candidate assignments rather than established causal genes. As such, the purpose of this analysis is to compare the association statistics obtained in probablistic and deterministic ATTICUS(+V) HET S-TWAS with those obtained in scPrediXcan using the same GWAS inputs, in terms of their signal specificity to previously identified GWAS hits. It is not to demonstrate the robustness or biological relevance of our findings - these are demonstrated in other evaluation analyses. In terms of the comparison with published scTWAS results [23], we note that the latter is affected by differences in GWAS power, gene coverage, cell-type definitions and significance thresholds used in the scTWAS analyses. Accordingly, hits absent from the published results are newly detected relative to that comparison, rather than necessarily representing previously unknown disease mechanisms. Finally, our sign concordance evaluation are performed using direction of effect of a gene's expression on SLE or AD in published Mendelian Randomization analyses [21, 22] (**Figure 3a**, **Appendix Figure 17**). The effect directions they have derived depend on the instrument validity, allele harmonisation and cell-type matching in their analyses, and do not necessarily establish causal expression-mediated effects.

#### G.4 Spurious or undiscovered S-TWAS genes

Identification of novel significant S-TWAS associations may arise through LD with genes with previously identified S-TWAS associations, or correlated predicted expression among genes and cell-types. To address the former we used LD blocks [17] to count distinct loci containing S-TWAS associations, alongside the number of significant genes, such that we do not over-report our model gains; this does not solve the problem. The latter cannot be adequately addressed within this paper. It requires the S2F backbone to be trained in a cell-type aware manner, using high quality single cell gene expression data, so that cell-type specific effects may be resolved from cell-type shared effects [25].

#### G.5 Cell-type specificity

Cell-type specificity in S-TWAS findings of statistical significance does not necessarily imply cell-type specificity of the underlying biological effect (**Appendix Figure 19**). Differences in cell abundance, expression measurement, prediction accuracy and statistical power can produce significance in one cell-type but not another. Our classifications of cell-type-specific and shared hits therefore describe the observed association patterns. CIGMA [25] computed cell-type specific and cell-type shared cis-heritability estimates provide complementary characterisation of cis-regulatory architecture, but analysis is restricted to overlapping cell-type definitions and genes with available estimates (**Appendix Figure 8**).

##### G.6 Uncertainty about direction

Our uncertainty estimates did not reliably indicate sign accuracy in personalised expression predictions or concordance of downstream S-TWAS effect signs with reference directions. Previous work showed that weak variant-effect predictions frequently changed sign across S2F models trained on different cross-validation folds, whereas the most prominent predictions were more directionally stable and consistent with observed eQTL effects [26]. Negative correlations between personalised predictions and observed expression also often became positive in another model replicate, suggesting that disagreement across S2F models can reveal directional uncertainty that our estimates did not capture due to a singular frozen S2F backbone replicate.

#### H Appendix Tables

**Table 1: Cell-type coverage of OneK1K and ROSMAP.** Donors indicates the number of individuals with available expression data for at least 3 genes in each cell-type. Coverage is the percentage of paired donors represented: 981 for OneK1K and 396 for ROSMAP. Cell-types are ordered by decreasing coverage within each dataset.

| Cell-type | Donors | Coverage (%) |
|---|---|---|
| **OneK1K (981 total donors)** | | |
| Central memory CD4-positive, alpha-beta T cell | 981 | 100.00 |
| Effector memory CD4-positive, alpha-beta T cell | 981 | 100.00 |
| Effector memory CD8-positive, alpha-beta T cell | 981 | 100.00 |
| Memory B cell | 981 | 100.00 |
| Naive thymus-derived CD4-positive, alpha-beta T cell | 981 | 100.00 |
| Natural killer cell | 981 | 100.00 |
| Regulatory T cell | 981 | 100.00 |
| Naive B cell | 980 | 99.90 |
| Naive thymus-derived CD8-positive, alpha-beta T cell | 980 | 99.90 |
| Central memory CD8-positive, alpha-beta T cell | 977 | 99.59 |
| Transitional stage B cell | 977 | 99.59 |
| Gamma-delta T cell | 975 | 99.39 |
| CD14-positive monocyte | 959 | 97.76 |
| CD16-negative, CD56-bright natural killer cell, human | 952 | 97.04 |
| CD4-positive, alpha-beta cytotoxic T cell | 946 | 96.43 |
| CD14-low, CD16-positive monocyte | 930 | 94.80 |
| Mucosal-associated invariant T cell | 900 | 91.74 |
| Conventional dendritic cell | 875 | 89.19 |
| Plasmablast | 795 | 81.04 |
| Hematopoietic precursor cell | 705 | 71.87 |
| Double negative thymocyte | 678 | 69.11 |
| Plasmacytoid dendritic cell | 639 | 65.14 |
| Platelet | 534 | 54.43 |
| CD4-positive, alpha-beta T cell | 438 | 44.65 |
| Innate lymphoid cell | 319 | 32.52 |
| CD8-positive, alpha-beta T cell | 216 | 22.02 |
| Erythrocyte | 157 | 16.00 |
| Dendritic cell | 142 | 14.48 |
| Peripheral blood mononuclear cell | 92 | 9.38 |

Continued on next page

Table 1 (continued)

| Cell-type | Donors | Coverage (%) |
|---|---|---|
| **ROSMAP (396 total donors)** | | |
| Upper-layer intratelencephalic | 392 | 98.99 |
| Oligodendrocyte | 388 | 97.98 |
| Deep-layer intratelencephalic | 379 | 95.71 |
| Astrocyte | 366 | 92.42 |
| MGE interneuron | 365 | 92.17 |
| CGE interneuron | 362 | 91.41 |
| Oligodendrocyte precursor | 360 | 90.91 |
| Microglia | 341 | 86.11 |
| Deep-layer corticothalamic and 6b | 252 | 63.64 |
| Miscellaneous | 150 | 37.88 |
| LAMP5-LHX6 and Chandelier | 150 | 37.88 |
| Vascular | 81 | 20.45 |
| Deep-layer near-projecting | 71 | 17.93 |
| Splatter | 47 | 11.87 |
| Hippocampal CA1-3 | 41 | 10.35 |
| Hippocampal dentate gyrus | 39 | 9.85 |
| Amygdala excitatory | 32 | 8.08 |
| Thalamic excitatory | 32 | 8.08 |
| Committed oligodendrocyte precursor | 24 | 6.06 |
| Fibroblast | 23 | 5.81 |
| Midbrain-derived inhibitory | 20 | 5.05 |
| Choroid plexus | 17 | 4.29 |
| Ependymal | 15 | 3.79 |
| Hippocampal CA4 | 11 | 2.78 |
| Lower rhombic lip | 11 | 2.78 |
| Upper rhombic lip | 11 | 2.78 |
| Medium spiny neuron | 10 | 2.53 |
| Mammillary body | 9 | 2.27 |
| Bergmann glia | 7 | 1.77 |
| Cerebellar inhibitory | 7 | 1.77 |
| Eccentric medium spiny neuron | 4 | 1.01 |

**Table 2: Detailed distillation results.** This table shows the Spearman correlations averaged across genes for each cell-type, showing distillation fidelity with the teacher. Only l-ctPred has been trained and evaluated on across-gene expression percentiles, while the other models used quantitative expression.

| Cell-type | l-ctPred | l-ctPred* | Deterministic | | | Probabilistic | | |
|---|---|---|---|---|---|---|---|---|
| | | | HET pure | HET | ADD | HET pure | HET | ADD |
| **OneK1K** | | | | | | | | |
| CD14-low, CD16-positive monocyte | 0.7988 | 0.7944 | 0.7748 | 0.5335 | 0.5237 | 0.7680 | 0.5286 | 0.5103 |
| CD14-positive monocyte | 0.7988 | 0.7933 | 0.7715 | 0.5344 | 0.5235 | 0.7686 | 0.5296 | 0.5101 |
| CD16-negative, CD56-bright natural killer cell, human | 0.7994 | 0.7935 | 0.7697 | 0.5336 | 0.5243 | 0.7708 | 0.5271 | 0.5117 |
| CD4-positive, alpha-beta cytotoxic T cell | 0.8008 | 0.7957 | 0.7729 | 0.5342 | 0.5239 | 0.7686 | 0.5321 | 0.5112 |
| CD4-positive, alpha-beta T cell | 0.7999 | 0.7915 | 0.7724 | 0.5333 | 0.5214 | 0.7670 | 0.5269 | 0.5088 |
| CD8-positive, alpha-beta T cell | 0.8013 | 0.7900 | 0.7753 | 0.5307 | 0.5216 | 0.7671 | 0.5292 | 0.5095 |
| Central memory CD4-positive, alpha-beta T cell | 0.8013 | 0.7945 | 0.7750 | 0.5377 | 0.5210 | 0.7668 | 0.5298 | 0.5081 |
| Central memory CD8-positive, alpha-beta T cell | 0.8017 | 0.7940 | 0.7719 | 0.5329 | 0.5231 | 0.7690 | 0.5296 | 0.5103 |
| Conventional dendritic cell | 0.7985 | 0.7942 | 0.7753 | 0.5363 | 0.5253 | 0.7671 | 0.5317 | 0.5120 |
| Dendritic cell | 0.7996 | 0.7933 | 0.7723 | 0.5366 | 0.5261 | 0.7677 | 0.5322 | 0.5124 |
| Double negative thymocyte | 0.8002 | 0.7927 | 0.7738 | 0.5365 | 0.5246 | 0.7688 | 0.5319 | 0.5119 |
| Effector memory CD4-positive, alpha-beta T cell | 0.8014 | 0.7931 | 0.7734 | 0.5327 | 0.5227 | 0.7669 | 0.5294 | 0.5102 |
| Effector memory CD8-positive, alpha-beta T cell | 0.7994 | 0.7938 | 0.7741 | 0.5330 | 0.5210 | 0.7674 | 0.5281 | 0.5083 |
| Erythrocyte | 0.8011 | 0.7953 | 0.7728 | 0.5332 | 0.5214 | 0.7663 | 0.5283 | 0.5082 |
| Gamma-delta T cell | 0.8006 | 0.7920 | 0.7752 | 0.5355 | 0.5236 | 0.7655 | 0.5305 | 0.5107 |
| Hematopoietic precursor cell | 0.8011 | 0.7928 | 0.7749 | 0.5364 | 0.5246 | 0.7664 | 0.5320 | 0.5121 |
| Innate lymphoid cell | 0.8016 | 0.7905 | 0.7745 | 0.5353 | 0.5231 | 0.7671 | 0.5307 | 0.5110 |
| Memory B cell | 0.8000 | 0.7915 | 0.7749 | 0.5334 | 0.5223 | 0.7689 | 0.5289 | 0.5090 |
| Mucosal-associated invariant T cell | 0.8004 | 0.7960 | 0.7749 | 0.5314 | 0.5238 | 0.7683 | 0.5289 | 0.5115 |
| Naive B cell | 0.8014 | 0.7931 | 0.7736 | 0.5315 | 0.5211 | 0.7674 | 0.5270 | 0.5072 |
| Naive thymus-derived CD4-positive, alpha-beta T cell | 0.8022 | 0.7932 | 0.7747 | 0.5345 | 0.5210 | 0.7693 | 0.5295 | 0.5081 |
| Naive thymus-derived CD8-positive, alpha-beta T cell | 0.8013 | 0.7926 | 0.7729 | 0.5331 | 0.5227 | 0.7678 | 0.5294 | 0.5101 |
| Natural killer cell | 0.7995 | 0.7917 | 0.7727 | 0.5325 | 0.5209 | 0.7681 | 0.5277 | 0.5075 |
| Peripheral blood mononuclear cell | 0.7987 | 0.7954 | 0.7731 | 0.5357 | 0.5213 | 0.7680 | 0.5293 | 0.5083 |
| Plasmablast | 0.8022 | 0.7965 | 0.7758 | 0.5344 | 0.5233 | 0.7702 | 0.5302 | 0.5103 |
| Plasmacytoid dendritic cell | 0.8003 | 0.7927 | 0.7747 | 0.5343 | 0.5229 | 0.7692 | 0.5300 | 0.5098 |
| Platelet | 0.8009 | 0.7937 | 0.7724 | 0.5314 | 0.5194 | 0.7663 | 0.5271 | 0.5076 |
| Regulatory T cell | 0.8013 | 0.7957 | 0.7735 | 0.5340 | 0.5221 | 0.7690 | 0.5293 | 0.5093 |
| Transitional stage B cell | 0.8011 | 0.7949 | 0.7776 | 0.5329 | 0.5223 | 0.7682 | 0.5284 | 0.5088 |
| Mean | 0.8005 | 0.7935 | 0.7738 | 0.5340 | 0.5227 | 0.7679 | 0.5294 | 0.5098 |
| SD | 0.0011 | 0.0016 | 0.0016 | 0.0018 | 0.0015 | 0.0012 | 0.0016 | 0.0015 |
| Minimum | 0.7985 | 0.7900 | 0.7697 | 0.5307 | 0.5194 | 0.7655 | 0.5269 | 0.5072 |
| Maximum | 0.8022 | 0.7965 | 0.7776 | 0.5377 | 0.5261 | 0.7708 | 0.5322 | 0.5124 |

Continued on next page

Table 2 (continued)

| Cell-type | l-ctPred | l-ctPred* | Deterministic | | | Probabilistic | | |
|---|---|---|---|---|---|---|---|---|
| | | | HET pure | HET | ADD | HET pure | HET | ADD |
| **ROSMAP** | | | | | | | | |
| Amygdala excitatory | 0.8033 | 0.7856 | 0.7631 | 0.5052 | 0.4915 | 0.7566 | 0.5009 | 0.4803 |
| Astrocyte | 0.8044 | 0.7900 | 0.7606 | 0.5086 | 0.4966 | 0.7543 | 0.5042 | 0.4843 |
| Bergmann glia | 0.8038 | 0.7898 | 0.7624 | 0.5124 | 0.4996 | 0.7558 | 0.5075 | 0.4874 |
| Cerebellar inhibitory | 0.8031 | 0.7882 | 0.7632 | 0.5256 | 0.5132 | 0.7559 | 0.5205 | 0.5000 |
| CGE interneuron | 0.8040 | 0.7881 | 0.7633 | 0.5029 | 0.4890 | 0.7547 | 0.4991 | 0.4786 |
| Choroid plexus | 0.8047 | 0.7890 | 0.7605 | 0.5002 | 0.4863 | 0.7542 | 0.4962 | 0.4756 |
| Committed oligodendrocyte precursor | 0.8039 | 0.7914 | 0.7614 | 0.5095 | 0.4949 | 0.7583 | 0.5056 | 0.4839 |
| Deep-layer corticothalamic and 6b | 0.8039 | 0.7874 | 0.7631 | 0.5125 | 0.4891 | 0.7586 | 0.5036 | 0.4783 |
| Deep-layer intratelencephalic | 0.8039 | 0.7883 | 0.7605 | 0.5028 | 0.4897 | 0.7534 | 0.4988 | 0.4785 |
| Deep-layer near-projecting | 0.8040 | 0.7909 | 0.7624 | 0.5064 | 0.4930 | 0.7544 | 0.5018 | 0.4815 |
| Eccentric medium spiny neuron | 0.8030 | 0.7883 | 0.7640 | 0.5013 | 0.4869 | 0.7561 | 0.4975 | 0.4766 |
| Ependymal | 0.8044 | 0.7878 | 0.7599 | 0.5078 | 0.4941 | 0.7537 | 0.5031 | 0.4829 |
| Fibroblast | 0.8043 | 0.7884 | 0.7618 | 0.5107 | 0.4980 | 0.7553 | 0.5061 | 0.4858 |
| Hippocampal CA1-3 | 0.8029 | 0.7891 | 0.7617 | 0.5019 | 0.4876 | 0.7520 | 0.4979 | 0.4772 |
| Hippocampal CA4 | 0.8027 | 0.7872 | 0.7607 | 0.5216 | 0.5090 | 0.7557 | 0.5169 | 0.4961 |
| Hippocampal dentate gyrus | 0.8029 | 0.7901 | 0.7597 | 0.5013 | 0.4875 | 0.7561 | 0.4974 | 0.4769 |
| LAMP5-LHX6 and Chandelier | 0.8037 | 0.7893 | 0.7672 | 0.5031 | 0.4895 | 0.7536 | 0.4992 | 0.4783 |
| Lower rhombic lip | 0.8019 | 0.7872 | 0.7607 | 0.5259 | 0.5140 | 0.7562 | 0.5211 | 0.5005 |
| Mammillary body | 0.8022 | 0.7902 | 0.7603 | 0.5166 | 0.5040 | 0.7524 | 0.5120 | 0.4916 |
| Medium spiny neuron | 0.8024 | 0.7890 | 0.7609 | 0.4982 | 0.4840 | 0.7560 | 0.4944 | 0.4736 |
| MGE interneuron | 0.8038 | 0.7922 | 0.7617 | 0.5037 | 0.4906 | 0.7529 | 0.4995 | 0.4788 |
| Microglia | 0.8020 | 0.7894 | 0.7646 | 0.5159 | 0.5052 | 0.7568 | 0.5105 | 0.4909 |
| Midbrain-derived inhibitory | 0.8029 | 0.7909 | 0.7621 | 0.5061 | 0.4933 | 0.7506 | 0.5017 | 0.4817 |
| Miscellaneous | 0.8027 | 0.7871 | 0.7620 | 0.5085 | 0.4962 | 0.7558 | 0.5040 | 0.4838 |
| Oligodendrocyte precursor | 0.8037 | 0.7888 | 0.7634 | 0.5027 | 0.4891 | 0.7569 | 0.4986 | 0.4783 |
| Oligodendrocyte | 0.8033 | 0.7894 | 0.7608 | 0.5135 | 0.5010 | 0.7564 | 0.5083 | 0.4883 |
| Splatter | 0.8039 | 0.7915 | 0.7604 | 0.5072 | 0.4929 | 0.7550 | 0.5031 | 0.4824 |
| Thalamic excitatory | 0.8031 | 0.7865 | 0.7612 | 0.5105 | 0.4966 | 0.7534 | 0.5065 | 0.4857 |
| Upper rhombic lip | 0.8034 | 0.7904 | 0.7620 | 0.5245 | 0.5127 | 0.7585 | 0.5196 | 0.4991 |
| Upper-layer intratelencephalic | 0.8033 | 0.7898 | 0.7625 | 0.5032 | 0.4905 | 0.7554 | 0.4991 | 0.4787 |
| Vascular | 0.8026 | 0.7912 | 0.7631 | 0.5136 | 0.5009 | 0.7562 | 0.5087 | 0.4886 |
| Mean | 0.8034 | 0.7891 | 0.7620 | 0.5092 | 0.4957 | 0.7552 | 0.5046 | 0.4840 |
| SD | 0.0007 | 0.0016 | 0.0016 | 0.0076 | 0.0083 | 0.0019 | 0.0073 | 0.0074 |
| Minimum | 0.8019 | 0.7856 | 0.7597 | 0.4982 | 0.4840 | 0.7506 | 0.4944 | 0.4736 |
| Maximum | 0.8047 | 0.7922 | 0.7672 | 0.5259 | 0.5140 | 0.7586 | 0.5211 | 0.5005 |

**Table 3:** Mean number of significant S-TWAS genes per cell type, in blood and brain. Percentage changes follow the direction indicated for each disease.

| | **SLE**: blood → brain | | | **AD**: brain → blood | | |
|---|---|---|---|---|---|---|
| Method | Blood | Brain | Change | Brain | Blood | Change |
| scPrediXcan | 130.1 | 102.7 | −21.1% | 108.3 | 118.7 | +9.6% |
| Deterministic (HET) | 115.2 | 93.9 | −18.5% | 100.1 | 112.2 | +12.1% |
| Probabilistic (HET) | 168.4 | 154.0 | −8.5% | 155.6 | 151.4 | −2.7% |

**Table 4: Cell-type matching for comparison with SLE and AD associations from scTWAS.** Our OneK1K and ROSMAP cell populations were mapped to the published SLE and AD groups from scTWAS, respectively. For each gene, cell-type-specific TWAS $p$-values were combined within the listed groups using ACAT. Groups containing a single cell-type retain that cell-type's original $p$-value. $n$ denotes the number of aggregated cell-types.

| ACAT group | $n$ | Cell-types combined |
|---|---|---|
| **SLE – OneK1K (12 published cell-types)** | | |
| B_Mem | 1 | Memory B cells |
| B_IN | 1 | Naive B cells |
| CD4_NC | 2 | Naive CD4 T cells; central-memory CD4 T cells |
| CD8_NC | 2 | Naive CD8 T cells; central-memory CD8 T cells |
| CD4_ET | 2 | Effector-memory CD4 T cells; cytotoxic CD4 T cells |
| CD8_ET | 1 | Effector-memory CD8 T cells |
| CD8_S100B | 1 | Mucosal-associated invariant T cells (MAIT) |
| NK | 1 | Natural killer cells |
| NK_R | 1 | CD16-negative, CD56-bright NK cells |
| Mono_C | 1 | CD14-positive monocytes |
| Mono_NC | 1 | CD14-low, CD16-positive monocytes |
| DC | 1 | Dendritic cells |
| **AD – ROSMAP (6 published cell-types)** | | |
| Astrocyte | 1 | Astrocyte |
| Microglia | 1 | Microglia |
| Oligodendrocytes | 1 | Oligodendrocyte |
| OPCs | 1 | Oligodendrocyte precursor |
| Excitatory_Neurons | 4 | Upper-layer intratelencephalic; deep-layer intratelencephalic; deep-layer corticothalamic and 6b; deep-layer near-projecting |
| Inhibitory_Neurons | 3 | MGE interneuron; CGE interneuron; LAMP5-LHX6 and Chandelier |

**Table 5:** Cell-type-specific genomic inflation factors $\lambda_{\mathrm{GC}}$ computed from the published scPrediXcan results. Estimates were calculated from unadjusted association $p$-values in supplementary Tables S4–S14 (type 2 diabetes, T2D) and S15–S43 (systemic lupus erythematosus, SLE). $N$ denotes the number of genes.

| Trait | Cell-type | $N$ | $\lambda_{\mathrm{GC}}$ |
|---|---|---|---|
| T2D | Acinar | 18,585 | 1.638 |
| T2D | Activated stellate | 18,624 | 1.623 |
| T2D | Alpha | 18,593 | 1.648 |
| T2D | Beta | 18,611 | 1.622 |
| T2D | Delta | 18,564 | 1.638 |
| T2D | Ductal | 18,614 | 1.607 |
| T2D | Endothelial | 18,613 | 1.598 |
| T2D | Epsilon | 18,566 | 1.663 |
| T2D | Gamma | 18,611 | 1.639 |
| T2D | Immune | 18,613 | 1.597 |
| T2D | Quiescent stellate | 18,617 | 1.636 |
| SLE | $CD4^+$ $\alpha\beta$ cytotoxic T | 17,185 | 1.437 |
| SLE | $CD4^+$ $\alpha\beta$ T | 17,050 | 1.437 |
| SLE | $CD8^+$ $\alpha\beta$ T | 17,414 | 1.447 |
| SLE | CD14-low $CD16^+$ monocyte | 17,761 | 1.312 |
| SLE | $CD14^+$ monocyte | 17,761 | 1.401 |
| SLE | CD16-negative CD56-bright NK | 17,660 | 1.414 |
| SLE | Central memory $CD4^+$ T | 17,825 | 1.421 |
| SLE | Central memory $CD8^+$ T | 17,267 | 1.411 |
| SLE | Conventional dendritic | 16,670 | 1.424 |
| SLE | Dendritic | 15,556 | 1.427 |
| SLE | Double-negative thymocyte | 17,592 | 1.449 |
| SLE | Effector memory $CD4^+$ T | 16,987 | 1.411 |
| SLE | Effector memory $CD8^+$ T | 17,311 | 1.440 |
| SLE | Erythrocyte | 16,309 | 1.441 |
| SLE | $\gamma\delta$ T | 17,118 | 1.441 |
| SLE | Hematopoietic precursor | 17,695 | 1.394 |
| SLE | Innate lymphoid | 17,206 | 1.419 |
| SLE | Memory B | 16,870 | 1.427 |
| SLE | Mucosal invariant T | 17,702 | 1.448 |
| SLE | Naive B | 16,865 | 1.415 |
| SLE | Naive thymus-derived $CD4^+$ T | 17,765 | 1.407 |
| SLE | Naive thymus-derived $CD8^+$ T | 17,699 | 1.426 |
| SLE | Natural killer | 17,781 | 1.441 |
| SLE | PBMC | 16,786 | 1.413 |
| SLE | Plasmablast | 17,642 | 1.461 |
| SLE | Plasmacytoid dendritic | 16,907 | 1.399 |
| SLE | Platelet | 17,071 | 1.418 |
| SLE | Regulatory T | 16,988 | 1.411 |
| SLE | Transitional-stage B | 17,756 | 1.415 |

**Table 6: ACAT-based cell-type grouping for UpSet plots.** For each gene, cell-type-specific TWAS $p$-values were combined within the listed groups using ACAT, yielding 12 groups from 29 OneK1K cell-types and 12 groups from 31 ROSMAP cell-types. Groups containing a single cell-type retain its original $p$-value. $n$ denotes the number of aggregated cell-types

| ACAT group | $n$ | Cell-types combined |
|---|---|---|
| **OneK1K (29 cell-types; 12 groups)** | | |
| Hematopoietic cell | 1 | Hematopoietic precursor cell |
| Plasmablast | 1 | Plasmablast |
| Platelet | 1 | Platelet |
| Erythrocyte | 1 | Erythrocyte |
| Lymphoid cell | 1 | Innate lymphoid cell |
| Mononuclear cell | 1 | Peripheral blood mononuclear cell |
| Thymocyte | 1 | Double-negative thymocyte |
| NK cell | 2 | Natural killer cell; CD16-negative, CD56-bright natural killer cell (human) |
| B cell | 3 | Memory B cell; naive B cell; transitional-stage B cell |
| Dendritic cell | 3 | Conventional dendritic cell; dendritic cell; plasmacytoid dendritic cell |
| Monocyte | 2 | CD14-low, CD16-positive monocyte; CD14-positive monocyte |
| T cell | 12 | CD4-positive $\alpha\beta$ T cell; CD4-positive $\alpha\beta$ cytotoxic T cell; CD8-positive $\alpha\beta$ T cell; central memory CD4-positive $\alpha\beta$ T cell; central memory CD8-positive $\alpha\beta$ T cell; effector memory CD4-positive $\alpha\beta$ T cell; effector memory CD8-positive $\alpha\beta$ T cell; $\gamma\delta$ T cell; mucosal-associated invariant T cell; naive thymus-derived CD4-positive $\alpha\beta$ T cell; naive thymus-derived CD8-positive $\alpha\beta$ T cell; regulatory T cell |
| **ROSMAP (31 cell-types; 12 groups)** | | |
| Astroglia | 2 | Astrocyte; Bergmann glia |
| Oligodendrocyte lineage | 3 | Committed oligodendrocyte precursor; oligodendrocyte; oligodendrocyte precursor |
| Microglia | 1 | Microglia |
| Vascular and stromal | 2 | Fibroblast; vascular |
| Choroid and ependymal | 2 | Choroid plexus; ependymal |
| Cortical excitatory | 4 | Deep-layer corticothalamic and 6b; deep-layer intratelencephalic; deep-layer near-projecting; upper-layer intratelencephalic |
| Limbic excitatory | 4 | Amygdala excitatory; hippocampal CA1–3; hippocampal CA4; hippocampal dentate gyrus |
| Subcortical excitatory | 2 | Mammillary body; thalamic excitatory |
| Inhibitory neurons | 5 | CGE interneuron; cerebellar inhibitory; LAMP5-LHX6 and Chandelier; MGE interneuron; midbrain-derived inhibitory |
| Medium spiny neurons | 2 | Eccentric medium spiny neuron; medium spiny neuron |
| Rhombic lip | 2 | Lower rhombic lip; upper rhombic lip |
| Other | 2 | Miscellaneous; Splatter |

### I Appendix Figures

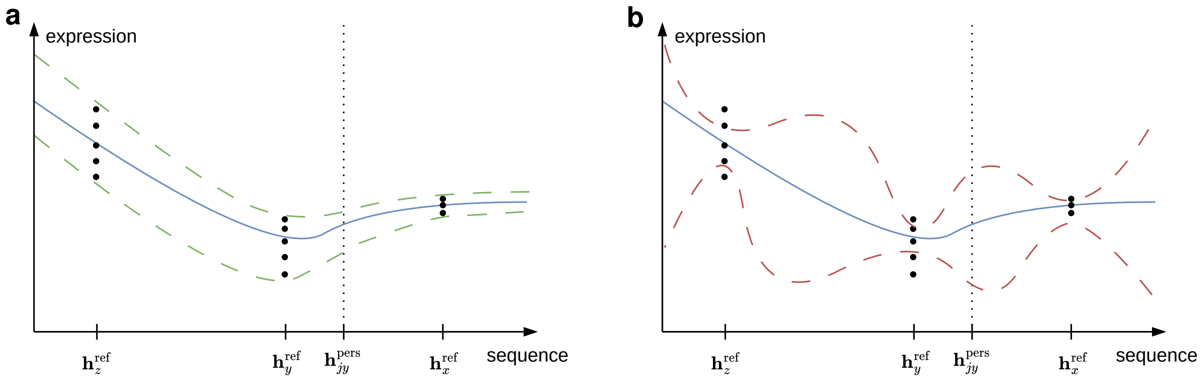


**Figure 1: Simplified illustration of training expression predictors in ATTICUS.** Each dot represents the pseudo-bulk expression of one individual for a fixed gene $g$ and cell-type $c$. The $y$-axis denotes expression magnitude, while the $x$-axis schematically represents the high-dimensional S2F embedding landscape. During training, expression measurements from different individuals are provided separately but are paired with the same reference-genome embedding $\mathbf{h}_g^{\text{ref}}$ for gene $g$. Across genes, this allows each ensemble member to learn a function mapping the reference embedding landscape to the conditional mean (blue), aleatoric variance (a, green) and epistemic uncertainty (b, red) of expression across individuals. At inference time, replacing the reference embedding with an individual-specific embedding $\mathbf{h}_{jg}^{\text{pers}}$ shifts the query within this learnt embedding landscape, which yields personalised estimates of the expression mean, variance and predictive uncertainty based on nearby reference gene representations seen during training.

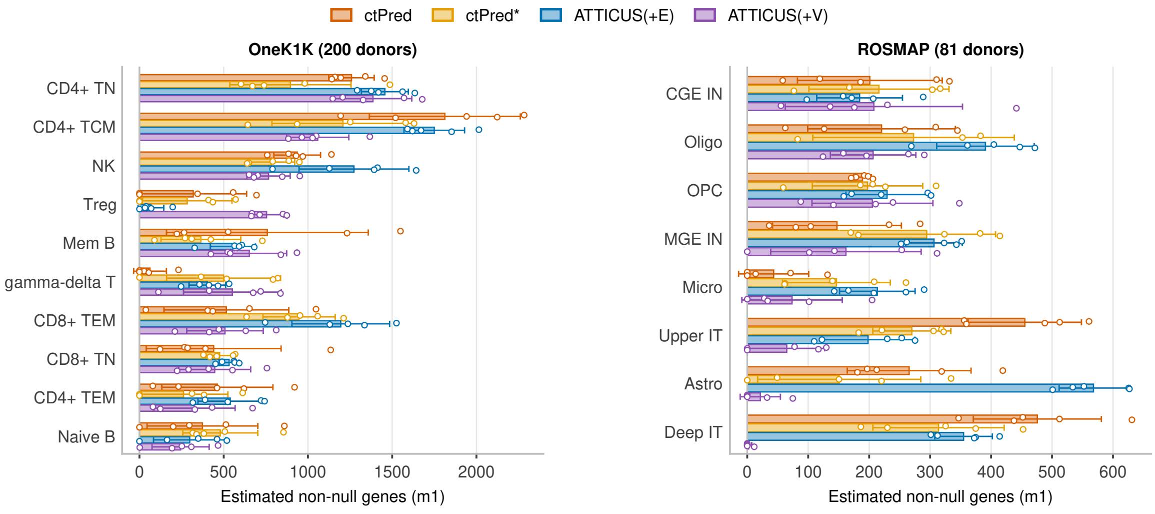


**Figure 2:** The $m_1$-metric is unstable across seeds for predictions for OneK1K and ROSMAP.

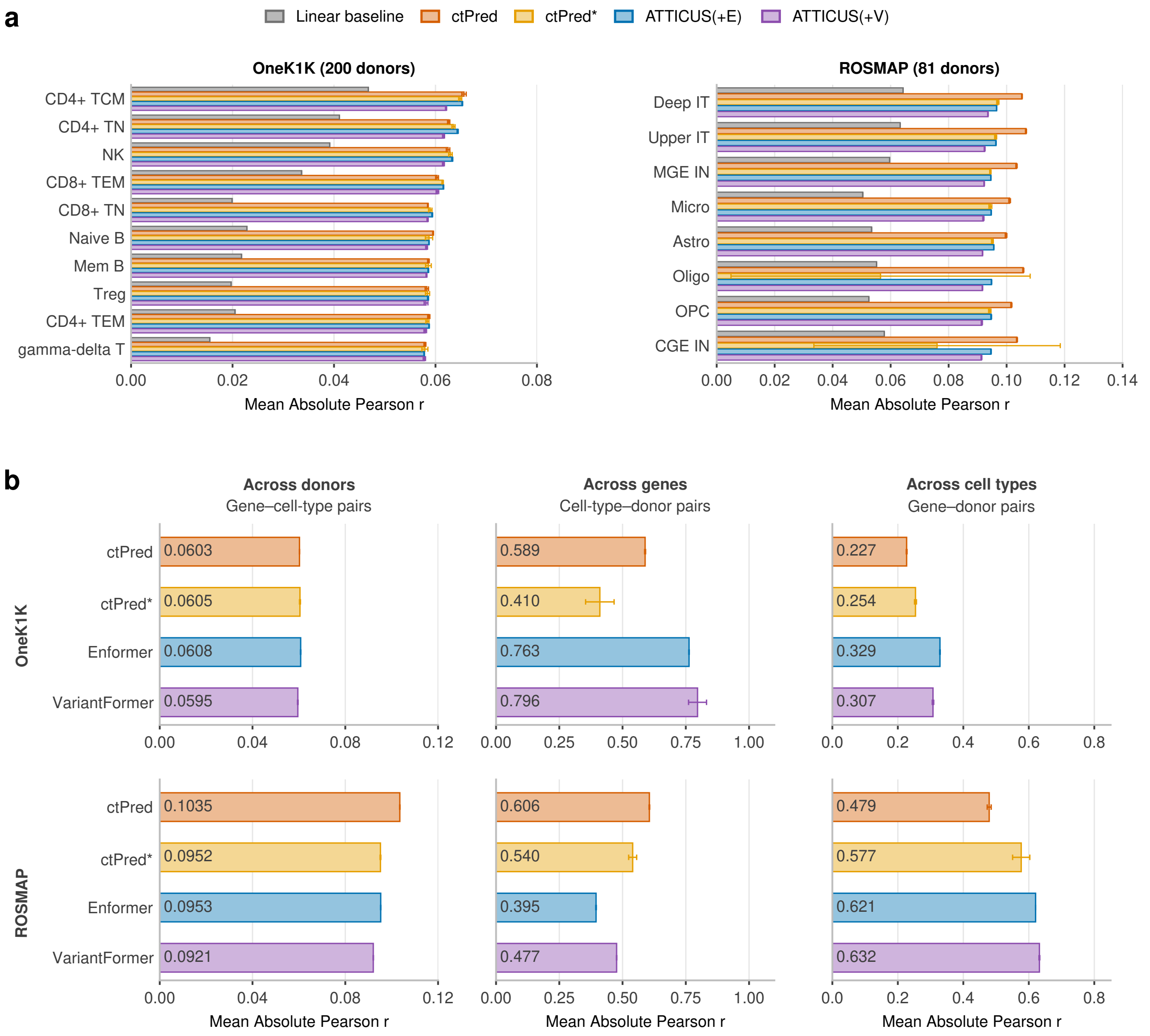


**Figure 3: Additional individualised expression prediction results.** a) All absolute expression prediction results, using all cell-types with full support. The linear baseline refers to a genotype-based elastic net trained directly on 80% of paired genotype–pseudo-bulk expression data. b) Individualised expression prediction results analysed across three different axes: across donors, across genes, and across cell-types.

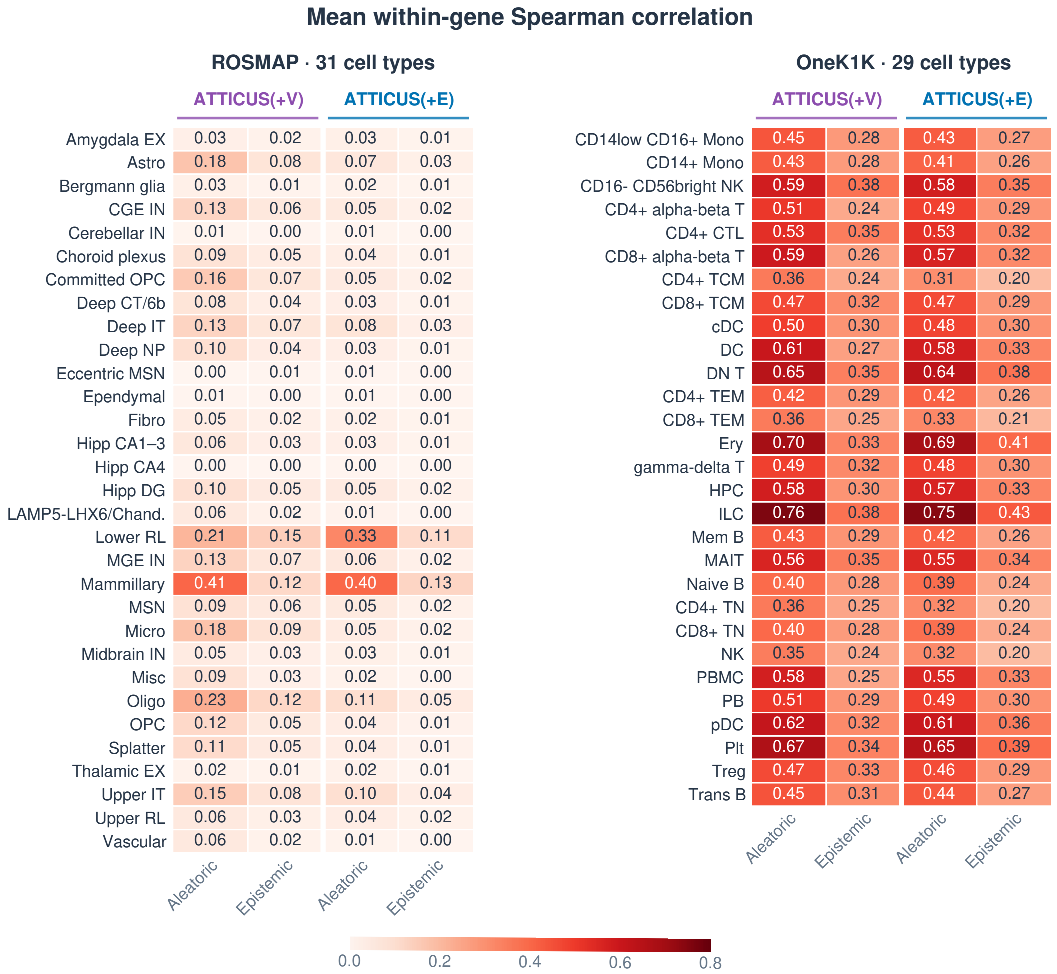


**Figure 4: Evaluating predictive uncertainty across individuals.** Mean within-gene Spearman correlation between predicted uncertainty and squared prediction error across individuals. Both metrics are evaluated separately for aleatoric and epistemic uncertainty, comparing ATTICUS(+E) and ATTICUS(+V) across all cell-types in OneK1K and ROSMAP.

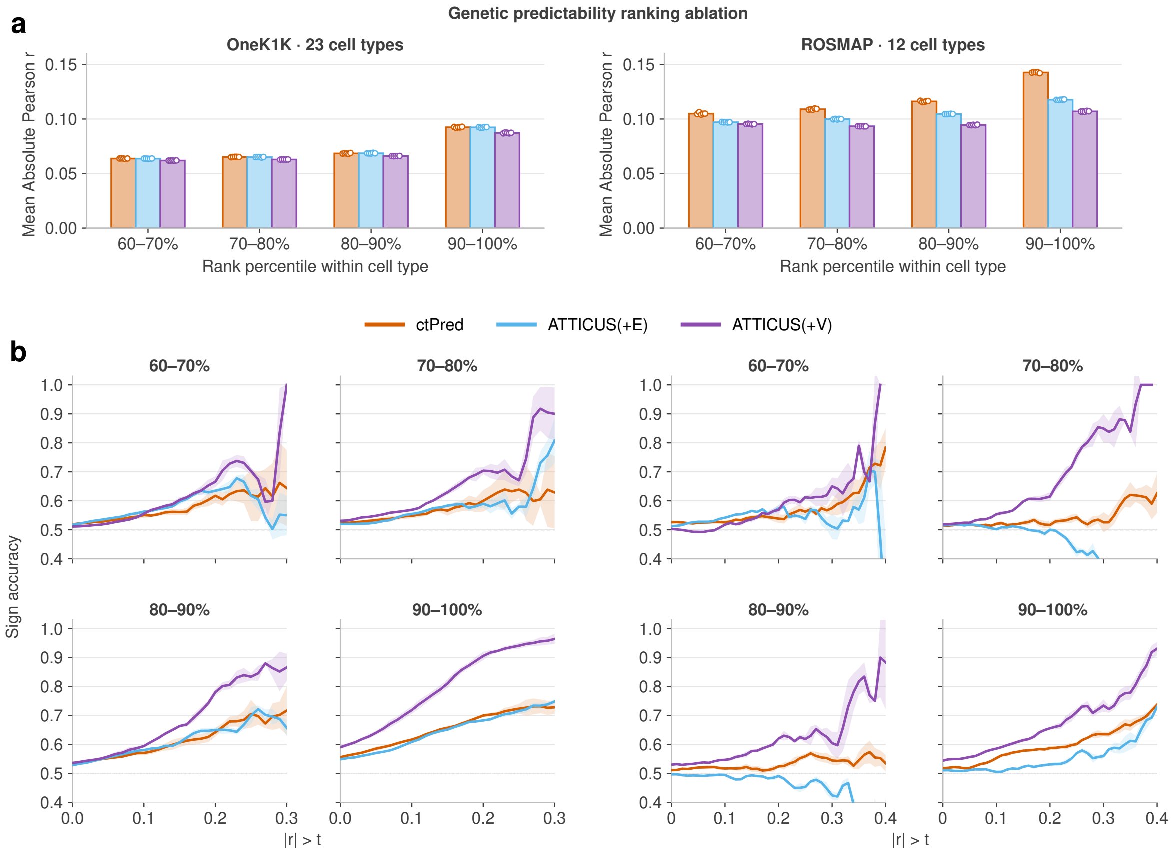


**Figure 5: Personalised prediction stratified by cis-predictability.** Genes were ranked within each cell-type by the held-out predictive performance of PrediXcan models trained on paired genotype–pseudobulk expression data. Prediction results are shown for the four highest deciles of predictability across 23 OneK1K and 12 ROSMAP cell-types with sufficient training data. Performance gains in both correlation magnitude and direction concordance are concentrated in genes with high cis-predictability. a) Mean absolute Pearson correlation between predicted and observed expression across donors, pooled across available cell-types. b) Sign accuracy across donors, pooled across available cell-types. The dashed line indicates the 50% chance level.

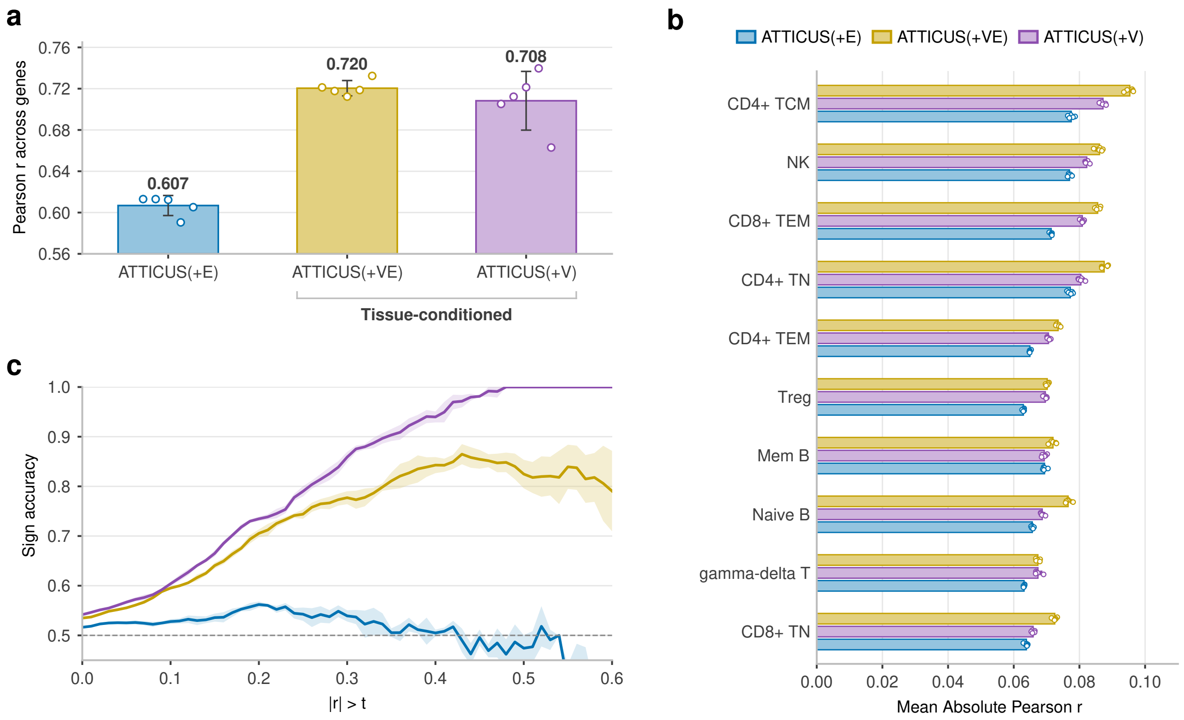


**Figure 6: Ablation of Variformer for personalised expression prediction.** a) Tissue-conditioned embeddings improve predictive performance on unseen genes on the reference. b) Individualised absolute prediction performance only shows little gain when variant-aware representations are used. c) While absolute predictive performance remains similar, sign accuracy (averaged across cell-types) increases substantially, especially for well-predicted genes.

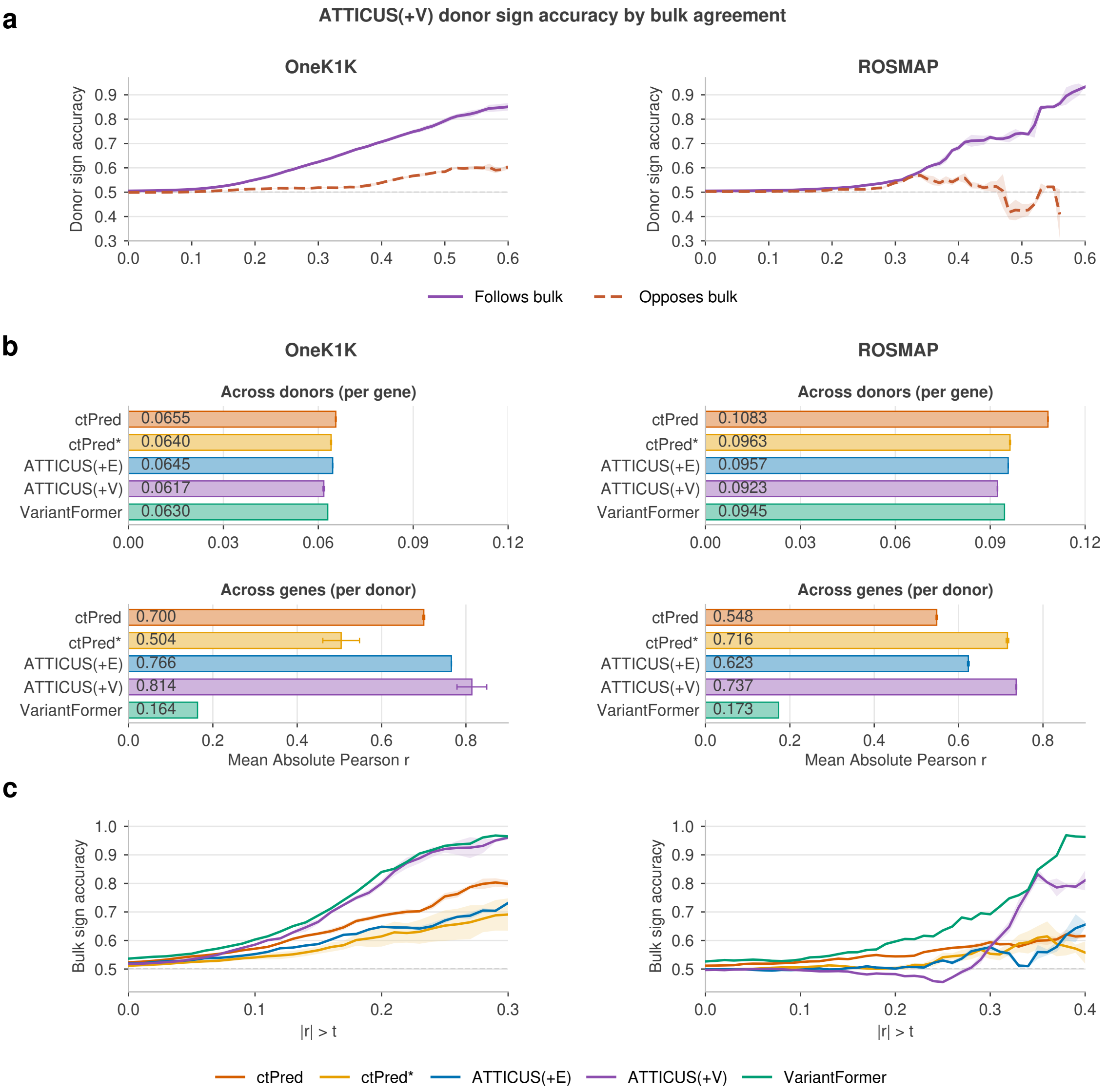


**Figure 7: Concordance of individualised predictions with bulk expression.** a) ATTICUS(+V) achieves higher sign accuracy for observed cell-type-specific pseudobulk expression when its predicted directions agree with zero-shot VariantFormer bulk predictions. This is consistent with transfer of bulk expression directions, while cell-type-specific directions (opposing the bulk prediction) remain harder to capture. b) Comparison of individualised bulk expression predictions from the evaluated models and zero-shot VariantFormer. The trained prediction heads substantially improve correlations across genes relative to zero-shot VariantFormer, whereas absolute correlations across donors remain similar. c) ATTICUS(+V) and zero-shot VariantFormer achieve the highest sign concordance with observed bulk expression, following similar trends across absolute-correlation thresholds.

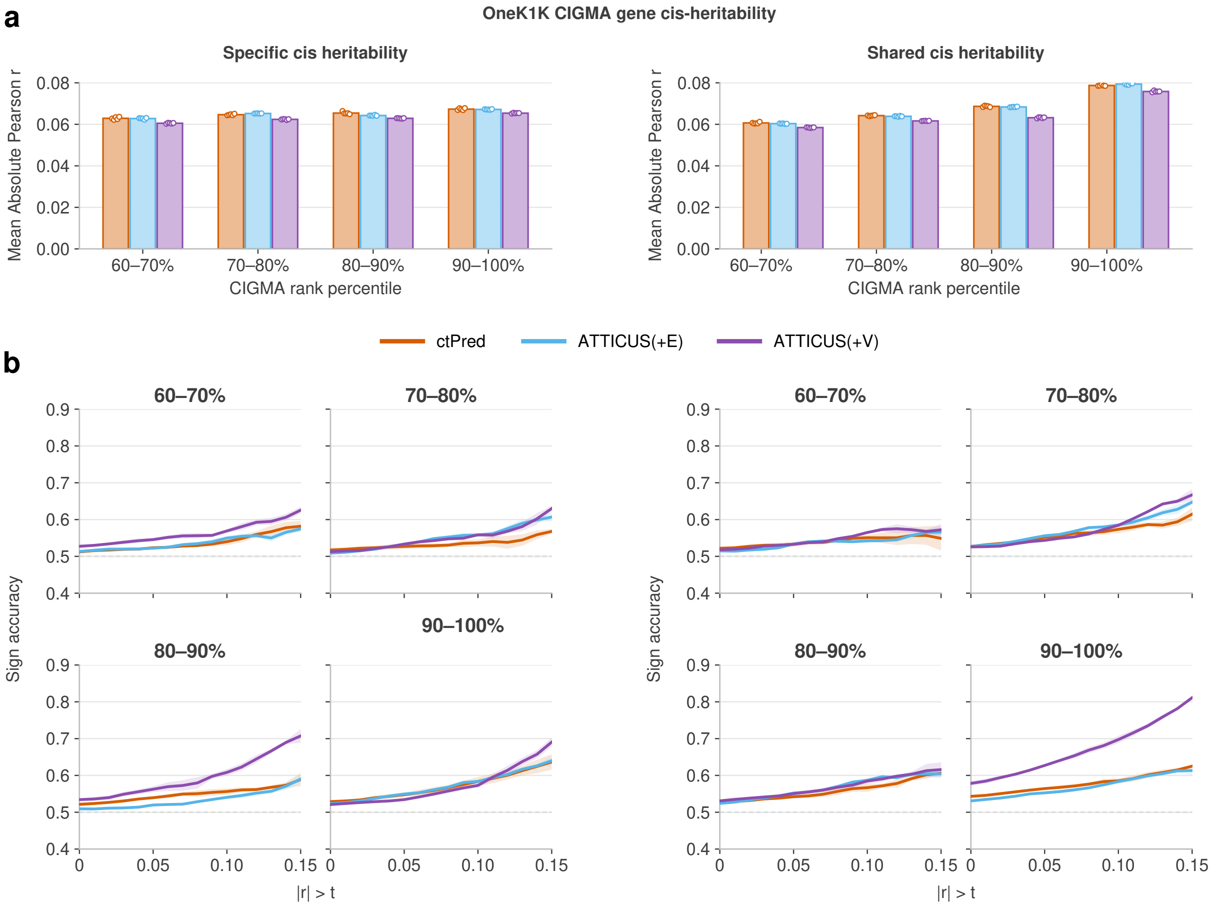


**Figure 8: Personalised prediction performance stratified by cell-type-specific and shared cis-heritability estimated by CIGMA.** Genes were ranked within seven available OneK1K cell-type populations by their CIGMA-estimated cell-type-specific or shared cis-heritability and grouped into the four highest deciles. Performance gains in both correlation magnitude and direction concordance are concentrated in genes with high shared cis-heritability. a) Mean absolute Pearson correlation between predicted and observed expression across held-out donors, pooled across available cell-types. b) Sign accuracy across donors, pooled across available cell-types. The dashed line indicated the 50% chance level.

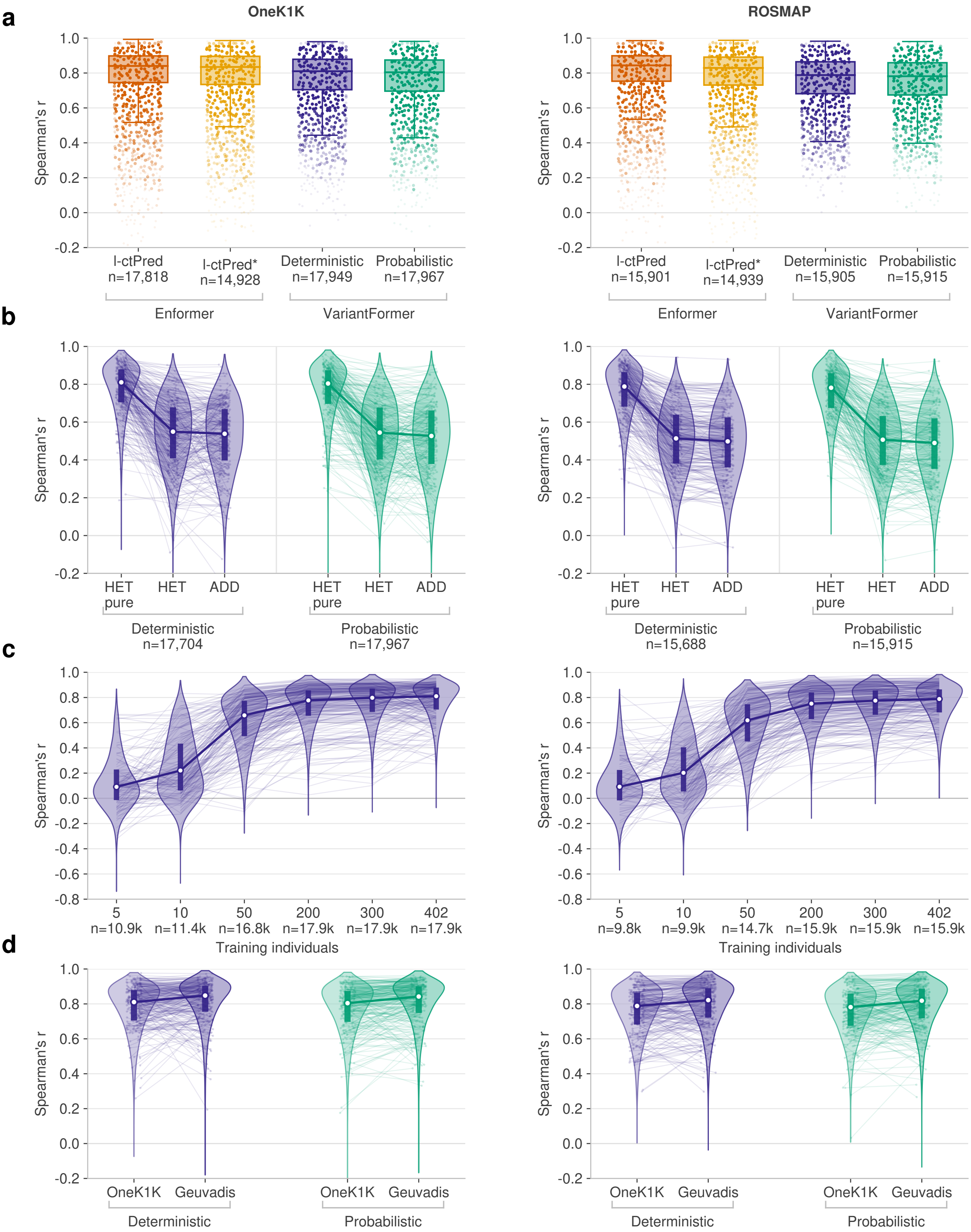


**Figure 9: Distillation fidelity across models and genotype panels.** Gene-wise teacher-student Spearman correlations across held-out individuals for OneK1K (left) and ROSMAP (right). a) Enformer-based l-ctPred and l-ctPred* compared with deterministic and probabilistic ATTICUS(+V) using full heterozygosity-aware students. b) Full heterozygosity-aware predictions (HETpure), their additive projection (HET) and directly fitted additive students (ADD). c) Effect of the number of individuals used for distillation training. d) Comparison of the original Geuvadis and OneK1K genotype panels, matched at 503 individuals each.

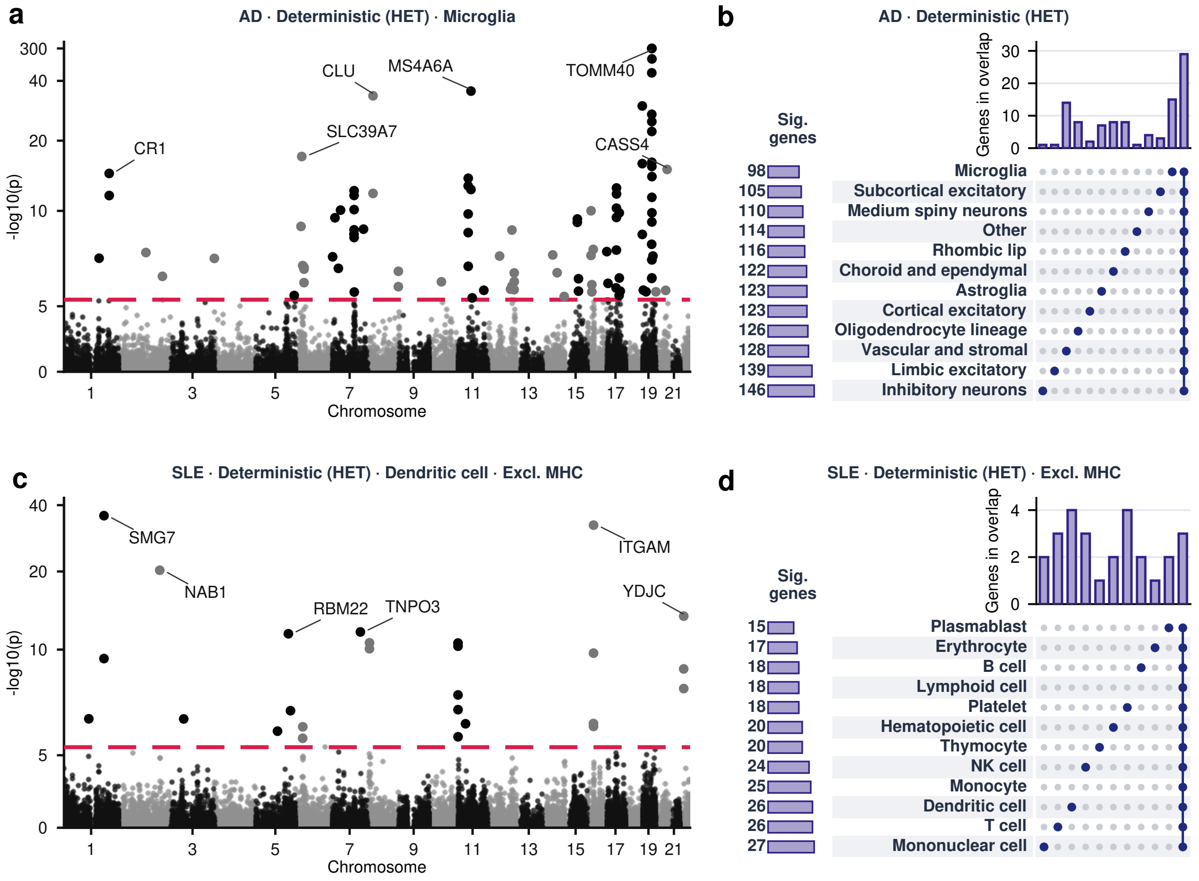


**Figure 10:** Cell-type-resolved TWAS associations identified by deterministic ATTICUS(+V) with heterozygote-aware distillation (HET) for SLE (blood) and AD (brain).

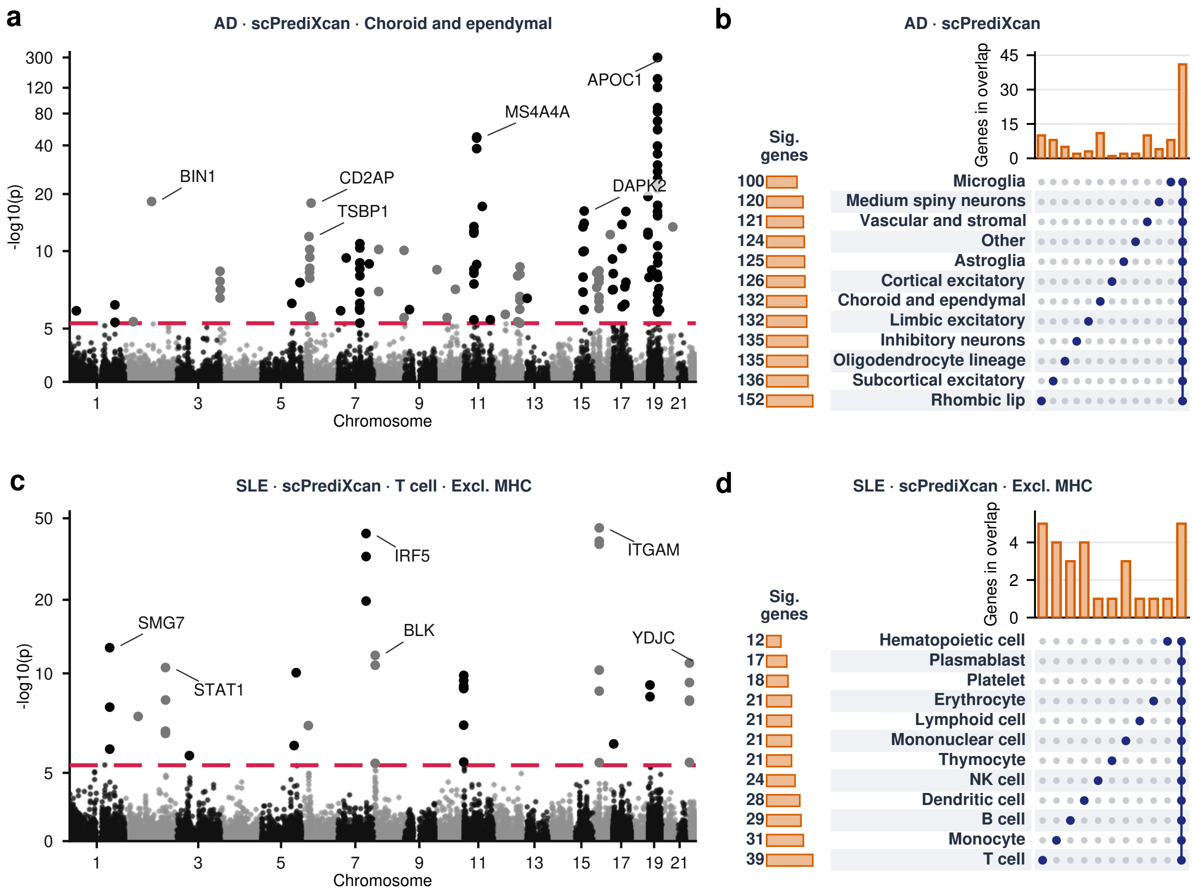


**Figure 11:** Cell-type-resolved TWAS associations identified by scPrediXcan for SLE (blood) and AD (brain).

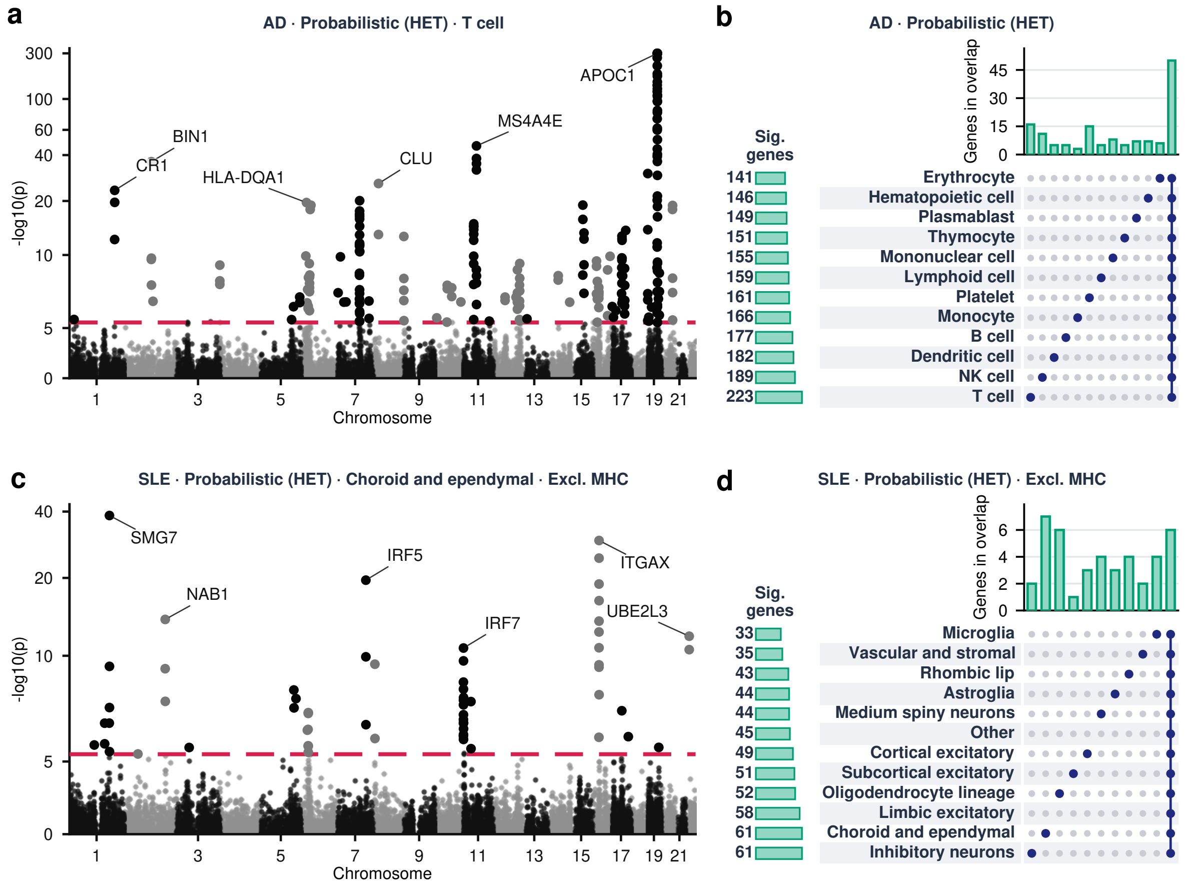


**Figure 12:** Cell-type-resolved TWAS associations identified by probabilistic ATTICUS (+V) with heterozygote-aware distillation (HET) for SLE (brain) and AD (blood).

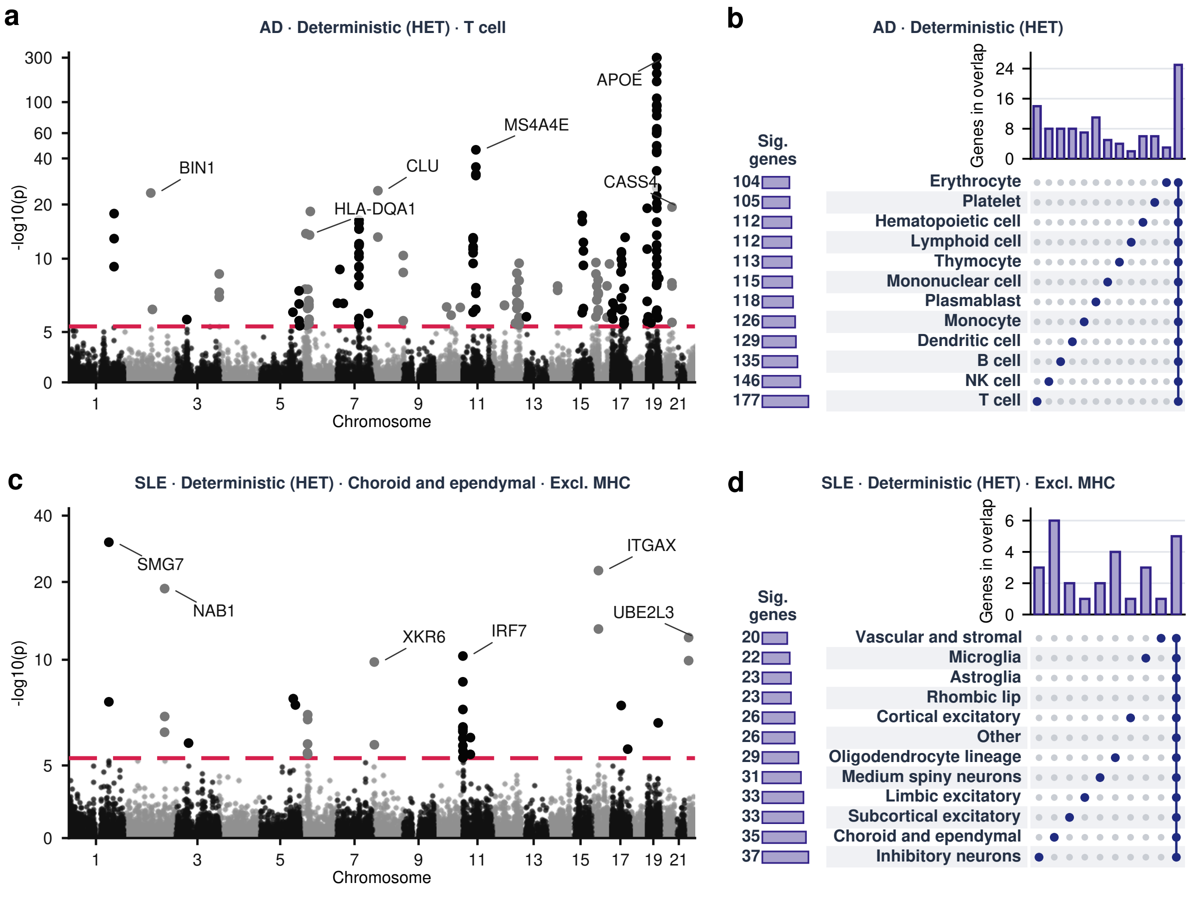


**Figure 13:** Cell-type-resolved TWAS associations identified by deterministic ATTICUS(+V) with heterozygote-aware distillation (HET) for SLE (brain) and AD (blood).

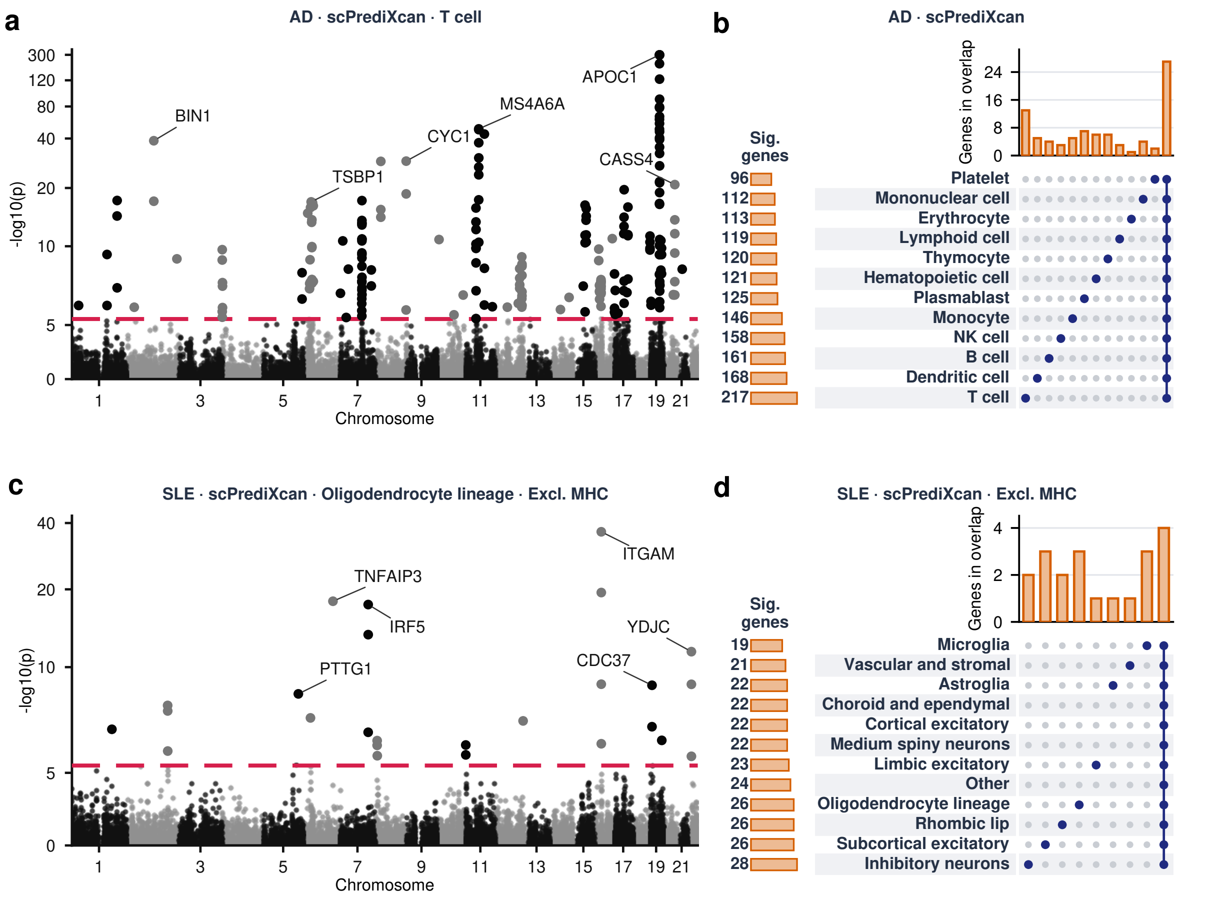


**Figure 14:** Cell-type-resolved TWAS associations identified by scPrediXcan for SLE (brain) and AD (blood).

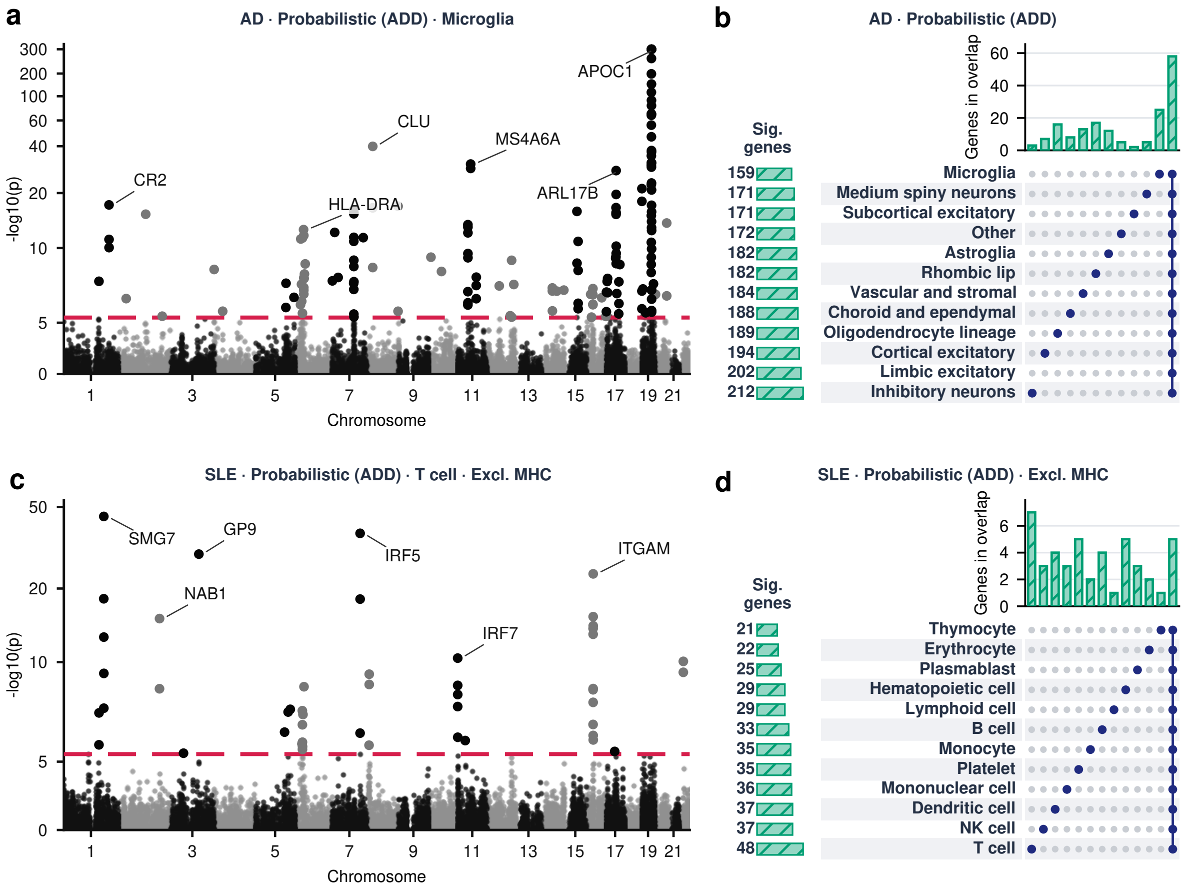


**Figure 15:** Cell-type-resolved TWAS associations identified by probabilistic ATTICUS (+V) with additive-only distillation (ADD) for SLE (blood) and AD (brain).

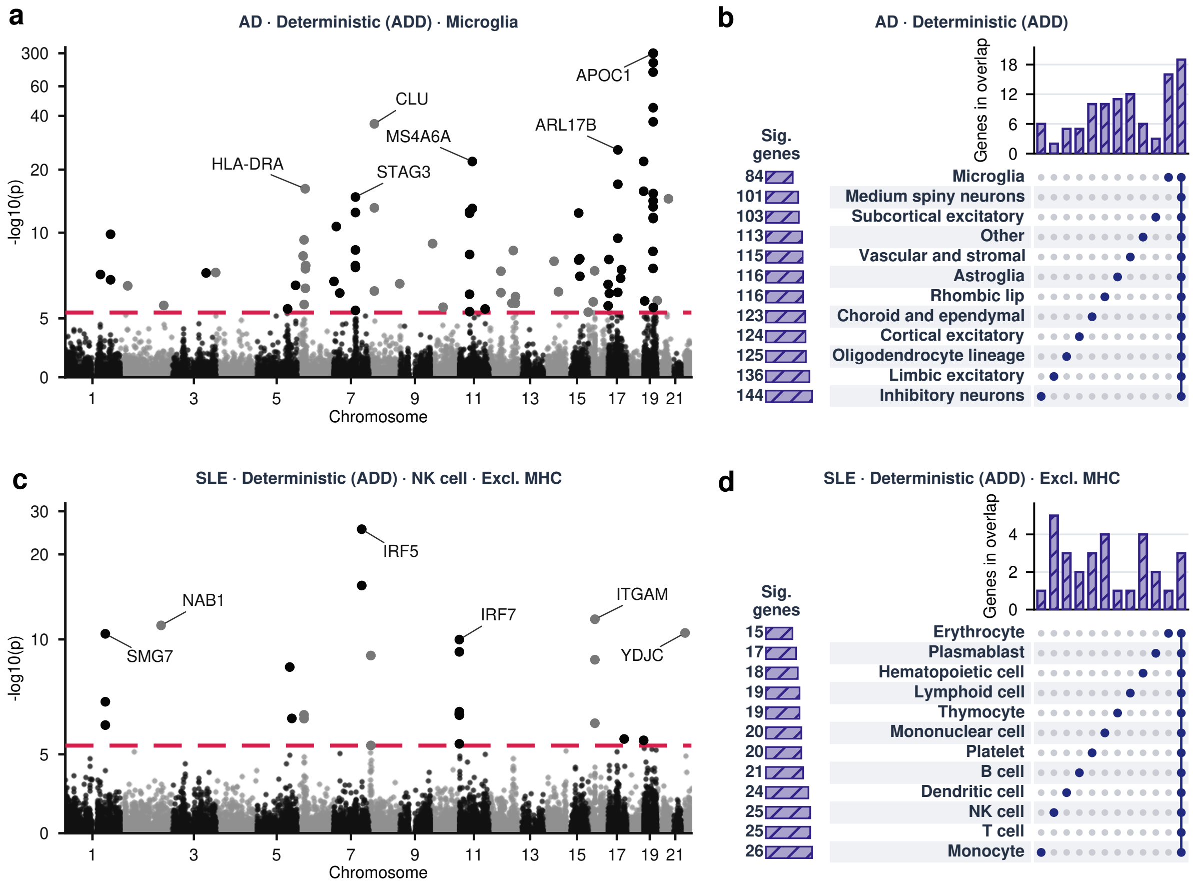


**Figure 16:** Cell-type-resolved TWAS associations identified by deterministic ATTICUS (+V) with additive-only distillation (ADD) for SLE (blood) and AD (brain).

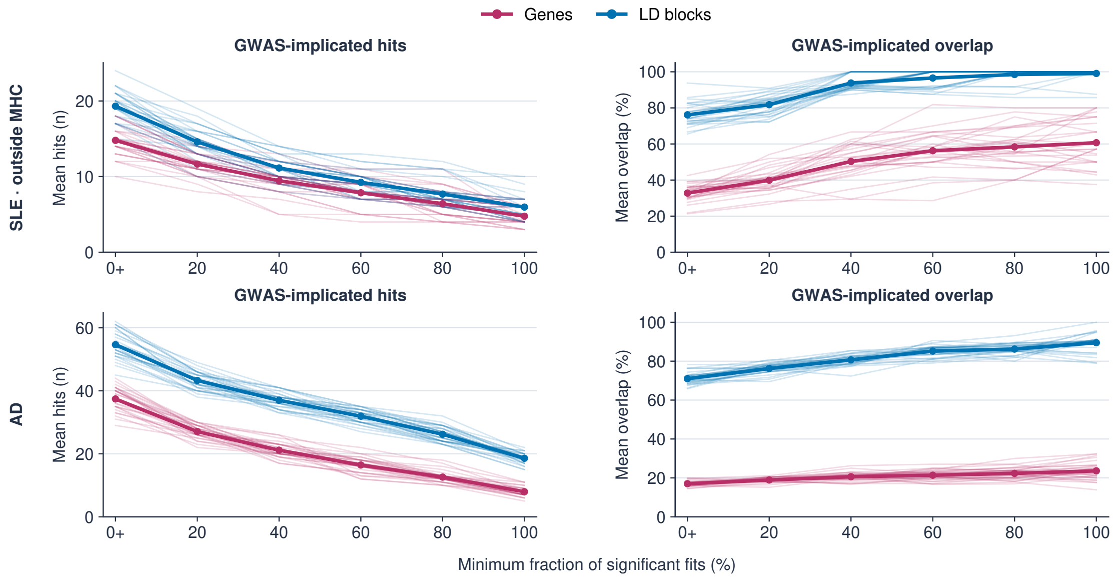


**Figure 17: Student-model agreement prioritises GWAS-supported TWAS associations.** Associations were kept only when the fraction of student models yielding Bonferroni-significant results met the threshold on the horizontal axis; 0+ denotes significance in at least one student model. Increasing the required agreement reduces the total number of retained GWAS-implicated hits (left) but increases their overlap with prior GWAS findings (right), which supports that student-model agreement is an informative measure of association uncertainty.

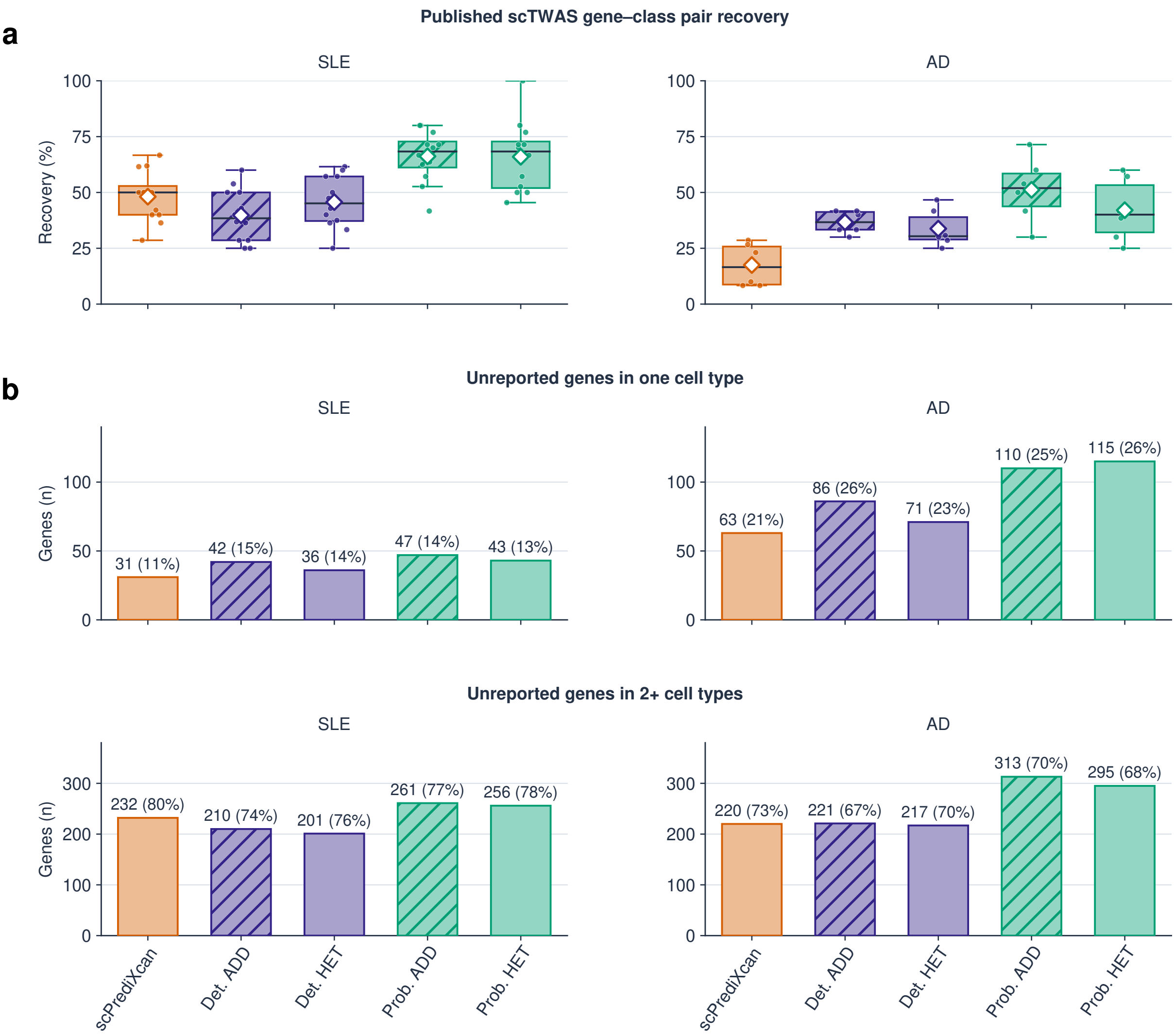


**Figure 18: Probabilistic TWAS improves recovery of published gene–cell-type associations and identifies additional hits detected in individual cell-types.** Comparison of scPrediXcan with deterministic and probabilistic ATTICUS for SLE and AD. a) Percentage of published scTWAS gene–cell-type associations recovered, summarised across matched cell-type groups. b) Number of additional genes not reported by scTWAS that reach significance in exactly one or at least two cell-type groups, respectively. Probabilistic ATTICUS generally recovers more previously reported associations and identifies more additional genes significant in only one group, particularly in AD. Proportions are comparable to those obtained with deterministic models, but numbers are further increased by their probabilistic counterparts.

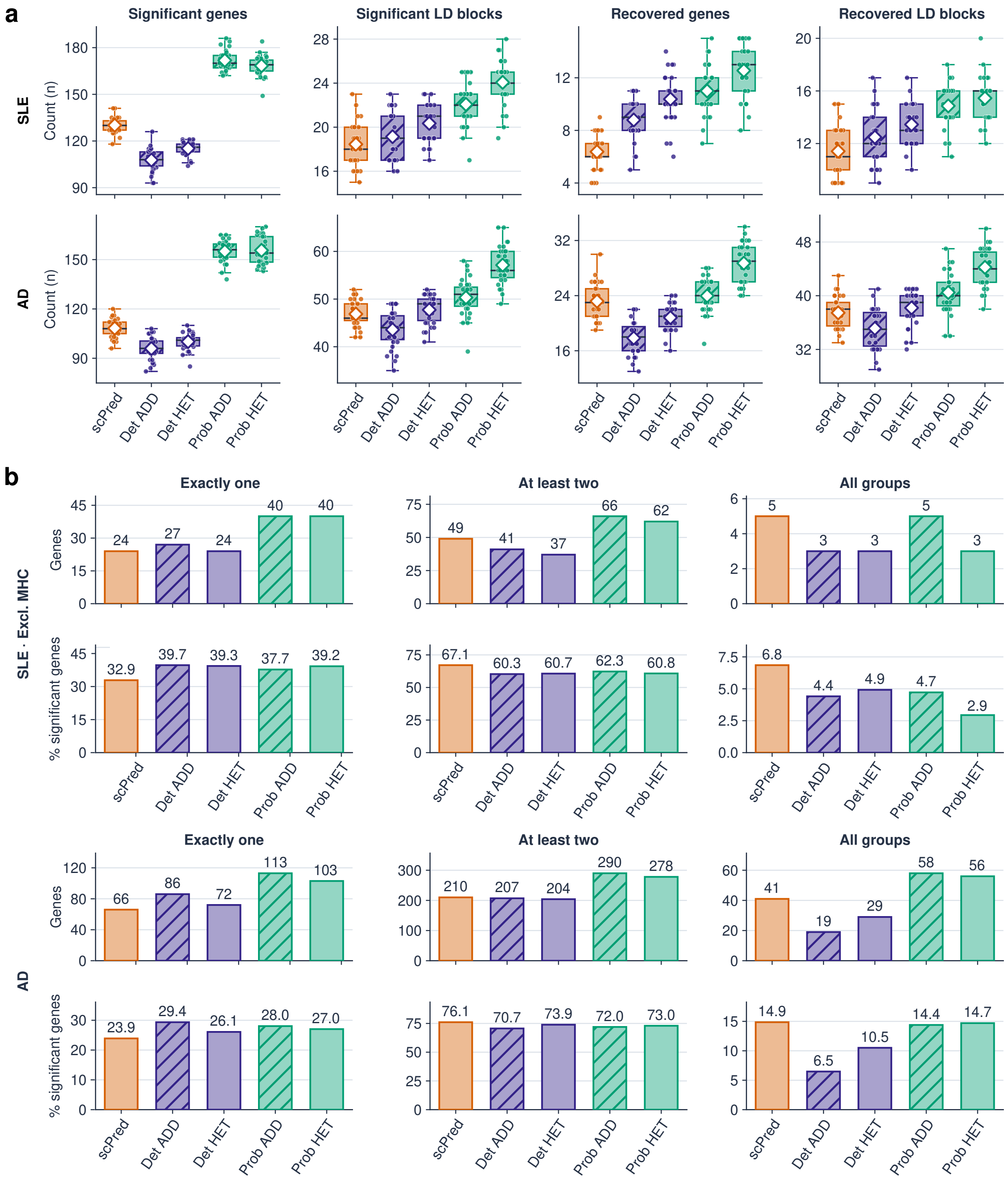


**Figure 19: TWAS discovery and cell-type specificity.** a) Number of significant genes and distinct LD blocks per cell-type, as well as number of recovered GWAS-implicated genes and LD blocks for SLE and AD. b) Number and proportion of significant genes identified in exactly one, at least two, or all major cell-type groups for SLE and AD.
